# Conformational changes upon pore blocker removal reveal conductive states of TMEM16A

**DOI:** 10.64898/2026.01.02.697443

**Authors:** Christina A. Stephens, Frank V. Marcoline, Christian J. Peters, Michael Grabe

**Author notes:** CORRESPONDANCE, (C.J.P.) or (M.G.). Irving Medical Center, Columbia University, New York, NY, 10032.

## Abstract

TMEM16A is a Ca^2+^-activated anion channel that provides direct electrical feedback to the plasma membrane in response to intracellular Ca^2+^. Its conductive state remains unresolved, leaving questions about gating, Cl^-^ permeation, and modulation by Ca^2+^, depolarization, and lipids. To investigate the open state, we performed molecular dynamics simulations of TMEM16A bound to the putative open-state blocker 1PBC. After inhibitor removal, the putative, pore-lining helix TM4 developed kinks at two sites: an upper site that opens the pore for Cl^-^ permeation, and a deeper site causing constriction. A conserved hydrophobic network between TM3 and TM4 persisted in most open structures but separated during extreme dilation, allowing lipids to transiently block the pore. Patch-clamp recordings indicated that the intact network promotes activation. Further simulations yielded >60 Cl^-^ permeation events and a single-channel conductance matching experiments. Additional electrostatic and kinetic modeling indicated that TMEM16A’s transition from outward-rectification to Ohmic conductance with increasing Ca^2+^ results from a weak voltage dependence of Ca^2+^ binding, which act cooperatively to open the pore.

**HIGHLIGHTS:**

- Removal of the inhibitor 1PBC from TMEM16A induces spontaneous transitions to ion conductive and non-conductive states via bending in TM4 at two different locations.
- Simulations suggest that the open state is stabilized by a small hydrophobic network between TM3 and 4 and disrupting this network biases channel closure in electrophysiological recordings.
- A kinetic model of conduction based on energetics from the all-atom MD simulations coupled to continuum calculations gives a linear current-voltage curve consistent with the fully open conformation and removal of 1 Ca^2+^ switches to an outwardly rectifying state via electrostatic influence on the Cl^-^ energy profile. However, the rectification is too weak to match experiment, but a cooperative model of Ca^2+^ binding with weak voltage-dependence does match experiment.

## INTRODUCTION

TMEM16A/Anoctamin1 is a Ca^2+^-activated Chloride Channel (CaCC)^1–3^ that is the archetype of the ten-member TMEM16 family of membrane-spanning proteins. Only TMEM16A and TMEM16B appear to behave exclusively as ion channels, while the other family members with known function act primarily as lipid scramblases. TMEM16A plays a role in a wide range of biological processes^4^ such as gastrointestinal motility^5^, fluid secretion^6^, nociception^7^, muscle contraction^8^, and regulation of blood pressure through tuning vascular tone^9^. It is also implicated in several cancers^10^, it can cause Moyamoya disease (a progressive cerebrovascular disorder)^11^, and it is a promising target for treating asthma and chronic obstructive pulmonary disease (COPD)^12^. In addition to Ca^2+^, TMEM16A conductance is modulated by phosphatidylinositol-4,5-bisphosphate (PI(4,5)P_2_) binding, which rescues current amplitudes following channel rundown with high, prolonged Ca^2+^ exposure^13–15^, and it is also modulated by voltage and extracellular anions^16,17^. Fatty acids, like arachidonic acid, are endogenous inhibitors of TMEM16A^18^, and they can regulate vascular tone as well^19^.

The structure of TMEM16A has been solved by cryo-electron microscopy (Cryo-EM) in apo, singly-bound, and doubly-bound Ca^2+^ states^20–22^. TMEM16A forms a homodimer with a butterfly-like fold, and each subunit has ten transmembrane (TM) segments and a hydrophilic cavity formed by TM3-7, which adopt an hourglass-like shape with the constriction point closer to the extracellular space. This domain forms the putative ion conduction pathway; however, none of the available TMEM16A structures have pores wide enough to accommodate a dehydrated Cl^-^ ion, making it unlikely that any of these structures reflects a conductive state of the protein (Table S1). Each subunit has two, adjacent Ca^2+^ binding sites made up of a cluster of acidic residues on TM6-8 on the cytoplasmic side. Ca^2+^ binding stabilizes an activated conformation of the Ca^2+^ binding domain enclosed by the intracellular ends of TM6-8 and induces more subtle rearrangements in the extracellular side of TM4^21,22^. While Ca^2+^ is activating, the existing doubly bound Ca^2+^ structures have been proposed to represent desensitized, non-conductive states due to the absence of PI(4,5)P_2_^22^. Meanwhile, the singly bound Ca^2+^ state may require positive voltage to open the pore, as TMEM16A conductance is voltage-dependent in low [Ca^2+^]^16^. In sub-saturating intracellular Ca^2+^ (270-400 nM), where the Ca^2+^-binding site may not be fully occupied, the channel is outwardly rectifying supporting inward Cl^-^ flow (positive current) under depolarizing voltages and very little flow at negative voltage^16^. As Ca^2+^ increases into and above low μM concentrations, the current-voltage relationship becomes linear (Ohmic), suggesting that TMEM16A may adopt more than one conductive state in different conditions. Unfortunately, the PI(4,5)P_2_ bound state has not been determined, and the structural and chemical details underpinning rectification and the Ca^2+^-dependent shift to Ohmic conductance remain unknown.

Important insight has come from all-atom molecular dynamics (MD) simulations that have focused on identifying the PI(4,5)P_2_ binding site(s) and how binding may open the pore^14,15,23^. MD simulations initiated from a Ca^2+^-bound structure (PDBID 5OYB^21^), after first docking PI(4,5)P_2_ into a putative binding site within TM3-5^23^, displayed spontaneous pore dilation wide enough to accommodate Cl^-^ ions and a single unbiased Cl^-^ permeation event; neither of these occurred in the absence of PI(4,5)P_2_ or Ca^2+^. Those authors suggested that pore opening primarily involves movement in TM3 and 4, and it relies on an allosteric network between the residues at the PIP_2_ site and TM4 residues on the extracellular side. Whether this conformation represents the Ohmic, outward rectifying, or some other state is not known, but calculations overestimated the selectivity of thiocyanate (SCN^-^)-to-Cl^-^ hinting that this state most resembles the outwardly rectifying state, which is more selective for SCN^-16^.

Experimental evidence also shows that TM3 is a dynamic element with some structures (PDBIDs 6BGI and 6BGJ^22^) exhibiting a ∼45° rotation of TM3 compared to others (PDBIDs 5OYG, 5OYB^21^, QZC^24^, 7B5E, 7B5D, and 7B5C^25^). In fact, channels adopt both TM3 positions independent of Ca^2+^, suggesting that the two states are visited at equilibrium^26^. This rotation moves the bulky residue Y514 out of the putative Cl^-^ permeation pathway replacing it with basic residue R515, which corroborates functional studies showing that mutation of R515, as well as pore lining residues K588, K603, and K645 on other TMs, modifies the conduction properties of the channel^16^. The structure of mTMEM16A has also been determined in the presence of the inhibitor 1PBC (PDBID 7ZK3^26^), and the molecule binds in the pore from the extracellular side capturing the rotation of TM3 that places R515 in the pore. 1PBC binding appears to bind the open state based on reduction in the IC_50_ with increasing Ca^2+^^16^ and more depolarized membrane potentials^26^.

Despite the availability of TMEM16A structures in multiple states, the lack of an open Cl^-^pore leaves many unanswered questions regarding the structural determinants of the Ohmic and rectifying conduction states and how the channel switches between them. We addressed these questions with all-atom MD starting from the 1PBC-bound structure. We hypothesized that if 1PBC blocks a conductive state of the channel, simulated removal of the inhibitor may reveal a conductive state that allows for Cl^-^ permeation. In the absence of 1PBC, the restrained protein did not permeate Cl^-^; however, in 12 independent, restraint-free simulations, half of the channels dilated wide enough for full Cl^-^ occupancy of the pore, allowing us to observe six complete permeation events. Using dimensionality reduction and clustering, we identified structural elements in TM3 and TM4 that appeared critical for open state stability, and we validated some of these features via patch clamp electrophysiology. Constant electric field simulations of representative conformations with dilated pores predicted a single channel conductance of 4.4 pS, consistent with experimental estimates. From these results, we constructed a mathematical model of the channel that recapitulates the Ohmic current-voltage (I-V) relationship under saturating Ca^2+^ and captures the switch to outward rectification as Ca^2+^ levels drop. These results provide a novel structural landscape to understand the progressive steps involved in Ca^2+^ and voltage-dependent activation of the TMEM16A channel.

## RESULTS

### The pore opens upon 1PBC removal

Prior to initiating our simulations, we modeled the three unresolved cytosolic loops in the 1PBC/Ca^2+^-bound cryo-EM structure (PDB ID 7ZK3, Fig. 1A). The loops preceding TM3 (15 residues long) and TM7 (13 residues long) are often missing in TMEM16 structures, likely due to their inherent flexibility, and we were particularly concerned with the placement of the TM2-3 loop owing to its implication in PIP_2_ binding and pore opening.

**Figure 1.**
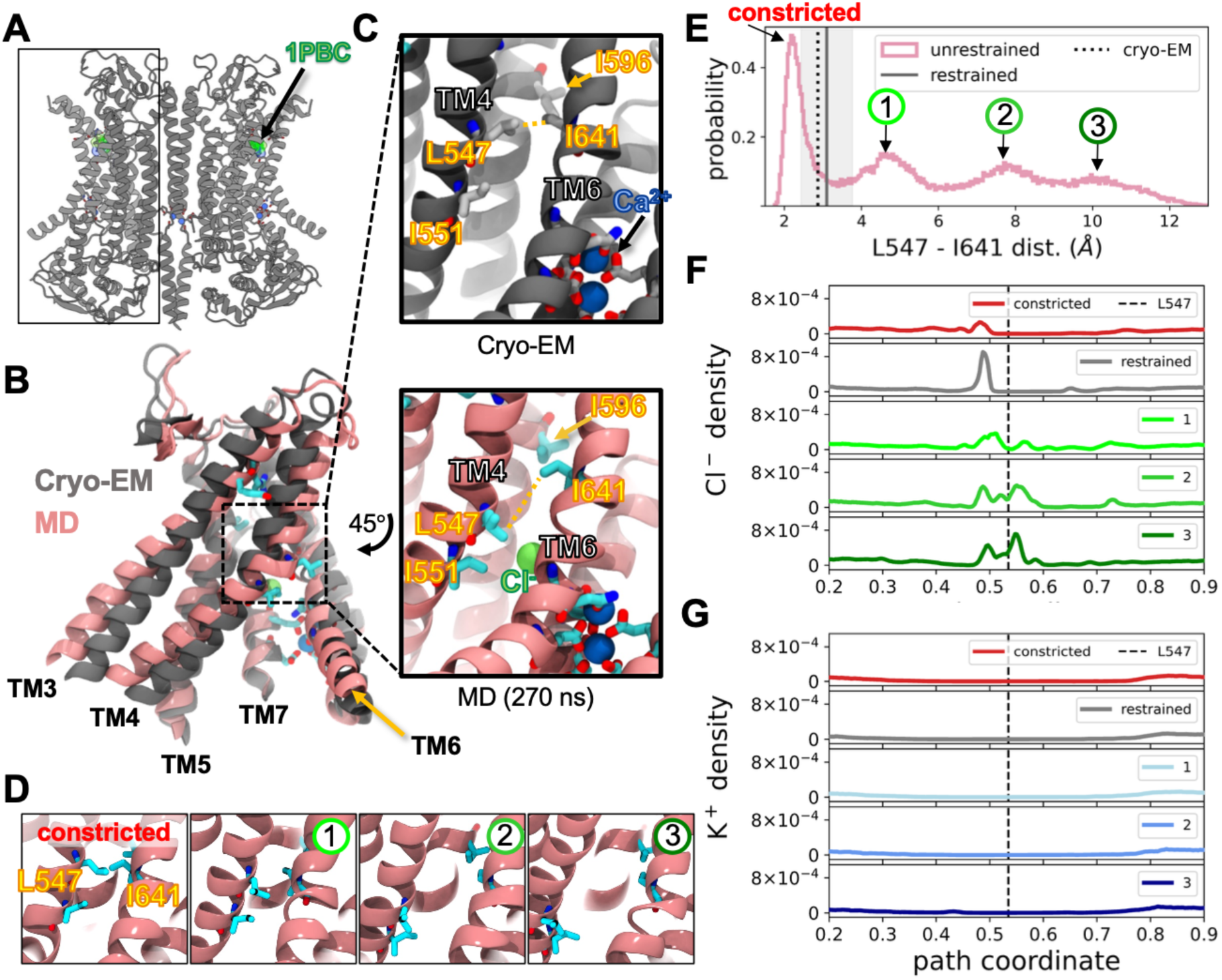
TMEM16A pore constricts or dilates the ion pathway when 1PBC is removed. **(A)** Cryo-EM structure of 1PBC (green)/Ca^2+^-bound (blue) TMEM16A (PDB ID 7ZK3). **(B)** Overlay of cryo-EM structure (gray) with snapshot from an unrestrained simulation (pink). **(C)** Enlarged view of Cl^-^ (green sphere) pathway constriction point (dashed box in panel B) for the structures in panel B. **(D)** Representative snapshots of 4 different degrees of gate separation. **I** Histogram of the minimum distances between gating residues L547 and I641 aggregated from all simulations. Vertical dotted line is the distance from cryo-EM structure, and the gray line and shaded region are the mean and standard deviation of distances from restrained simulation, respectively. Labeled peaks correspond to labels over snapshots in D **(F, G)** Average Cl^-^ and K^+^ densities along the pore separated into 4 categories based on the L547-I641 distance: 0-3 Å (constricted), 3-6 Å (1), 6-9 Å (2), and 9-20 Å (3). Densities from the restrained simulation are gray. These panels share the same x axis. Vertical dashed line is the position of L547 in the pore.

We generated four different models using MODELLER^27^ and AlphaFold2^28^ (Fig. S1): one with all loops built independently with MODELLER, one where we symmetrized the MODELLER-built loops between subunits, one where the AlphaFold2-built loops where also constructed symmetrically across subunits, and one where the cytosolic loops were omitted. Given the variation in loop positions among these models, we used all four in subsequent simulations.

Next, we tested the stability of the 1PBC-inhibited state after removal of the small molecule. We simulated three replicates of each of the four models for 1 μs, resulting in an aggregate simulation time of 12 μs per dimer, or 24 μs for all individual subunits. In each starting structure, the pore diameter is narrowest at a hydrophobic gate formed by residues on TM4 and TM6^29^. Within the first 300 ns of simulation, we observed that 12 out of 24 subunits spontaneously underwent major conformational changes causing the central part of the pore to widen (Fig. 1B, C). To quantify the changes in pore size, we measured the minimum distance between all atoms at hydrophobic gate residues L547 (TM4) and I641 (TM6). The distribution of aggregated distances revealed a multimodal landscape with peaks around 2.5, 5, 8, and 10 Å indicating that the simulations had sampled multiple metastable conformations (Fig. 1D-E). The smallest of the dilated peaks (peak 1) is characterized by a slight rotation and concomitant bend of TM4 at L547 exemplified by the snapshot shown in in Fig. 1B-C (pink). Here, L547 moves out of the pore, disengaging the hydrophobic gate as described in reference^23^ and the change in the helix is similar to a previously reported activating transition in which TM4 locally undergoes an α-to-π helix transition^30^. The peaks at the two larger distances (peaks 2 and 3 in Fig. 1E) involve larger-scale motions of TM4, including tilting into the membrane across TM6. Some pores collapsed further, causing greater constriction at L547-I641 (distances around 2.5 Å) than what is observed in the starting cryo-EM structure (vertical dotted line). This distance resembles values captured in Ca^2+^-free TMEM16A structures (Table S1), despite having Ca^2+^ bound throughout the simulations. All models sampled the constricted distribution and the dilated peak 2; how well they sampled each state varied between loop models. For example, simulations with symmetric MODELLER-based loops more frequently sampled distances in the 4.5-5 Å range (peak 1), while the asymmetric MODELLER-based loop model spent little time sampling this peak (Fig. S2).

We next looked at how these pore conformational changes altered ion accessibility to the pore. We measured the total density of Cl^-^ or K^+^ along a 3D path coordinate centered on the pore (see Methods and Fig. S3) from aggregate simulation data with L547 and I641 distances around the states indicated in Fig. 1E. In all conformations, including those from simulations restrained around the starting structure, the intracellular vestibule of the pore is accessible to Cl^-^(peak at 0.49 in Fig. 1F). For both the constricted and restrained starting conformations the density of Cl^-^ was near zero in the rest of the pore and likely not able to conduct ions. Despite separating the gating residues, we did not observe Cl^-^ ions in the middle of the pore after the first dilation event (1), suggesting the pore was not yet fully open. It was not until the gate separated farther (2 and 3 in Fig. 1F) that we saw Cl^-^ ions throughout the pore with a notable new peak above the L547 starting position (Fig. 1F). In all simulations the density of K^+^ was nearly zero in the pore which is consistent with experimental evidence that TMEM16A is anion selective (Fig. 1G)^17^.

To test if these observed conformational changes were due to the removal of 1PBC, we also initiated simulations with the inhibitor still bound. It is not known if 1PBC binds in its neutral or negativity charged, deprotonated state. We therefore simulated both states of the inhibitor initiated from the same cryo-EM pose in three, 1 μs replicates of each. When either the charged (1PBC(-1)) or neutral (1PBC(0)) was bound, TM4 moved away from its starting coordinates; however, these motions were not as severe as those we observed in the absence of inhibitor. In both cases the overall Cα RMSD of the extracellular-facing half of the pore to the starting structure was lower than in most simulations without 1PBC (Figs. S4I-J, S5). Despite subtle changes to the pore including separation of the hydrophobic gating residues (Fig. S4G, H), both forms of the inhibitor remained bound in the pore throughout the simulation, and we observed no ion permeation events. Of the two, 1PBC(-1) seemed to be the less stable form as it disengaged twice from the original binding location, moved deeper into the pore (Fig. S4B, right) and caused dilation (Fig. S4G). Given its weak binding affinity (∼3.6 μM K_d_ at 0 mV), it was not surprising that 1PBC would move around in the pore^26^; however, the 1PBC(0) form shifted only slightly in the original binding site bringing the protonated oxygen closer to E633 on TM6 (Fig. S4F).

### Two hinges on TM4 control opening

Pore opening correlates with a bend in TM4 at L547 resulting in a π-bulge in the helix, which compromises the integrity of the hydrophobic gate (Fig. S6A, D). Separately, closed structures also exhibit a bend in TM4, but at a different location centered on E555, two helical turns closer to the cytoplasm (Fig. S6B, E). Bending at E555 is accompanied by a movement of the extracellular terminus of TM4, which further occludes the extracellular half of the pore. We only observed kinking at both sites concurrently in one simulation derived from a model that lacked cytosolic loops. Additionally, we rarely observed TM4 adopt a fully straight conformation as it appears in the cryo-EM structure. Notably, neither location is centered on a glycine or proline residue.

We also observed clear changes in the number of waters in the center of the pore as a function of the helix bend angle at L547 or E555. When TM4 bends at L547 (to ∼160°), the number of waters occupying the center of the pore increases, whereas when it bends at E555 (to ∼150°), the pore dehydrates (Fig. S6A-E, G). When we applied restraints to maintain the starting conformation with a relatively straight TM4, there was only an intermediate amount of water (0-3 molecules) in the pore (Fig. S6C, F). The bend at L547 is also part of the TM4 motion that separates the hydrophobic gate, as indicated by increasing distances between L547-I641 in states corresponding to peaks 1, 2, and 3 (Figs. 1E, S8H). In contrast, the kink at E555 brings the hydrophobic gate closer together, as seen in the constricted states.

### Dilated states of TMEM16A are ion conductive

The inhibitor-free simulations revealed five outward permeation events and one inward event as seen in the time series traces through the channel (Fig. 2A and Fig. S7). All permeation events occurred when the L547-I641 distance was 9 Å or greater. The permeating ions dwell in three locations (sites A (outer), B (central), and C (inner)), which correspond to the three Cl^-^ density peaks in Fig. 1E. Site A is the least occupied according to the density profile and overlaps with the 1PBC binding site (Fig. S4A). Cl^-^ ions are simultaneously coordinated by R515 and K603 at the top of the extracellular vestibule (Fig. 2B, top), mimicking the 1PBC unprotonated oxygen binding pose. Site C is the second most occupied location and is formed by K645 (TM6) and K588 (TM5) in the cytosolic vestibule (Fig. 2B, bottom). There is also a third site, B, situated between A and C above the pore constriction site where the ion is coordinated by R515 and K645 (Fig. 2C). Interestingly, the middle site is formed by one residue from site A (R515) and one from site B (K645) in a “hand-off” scheme as the anion moves from one site to the next. Thus, the Cl^-^ binding sites in TMEM16A are somewhat fluid as long, flexible residues snorkel to coordinate ions at multiple locations, which has been observed in simulations by others^23^.

**Figure 2.**
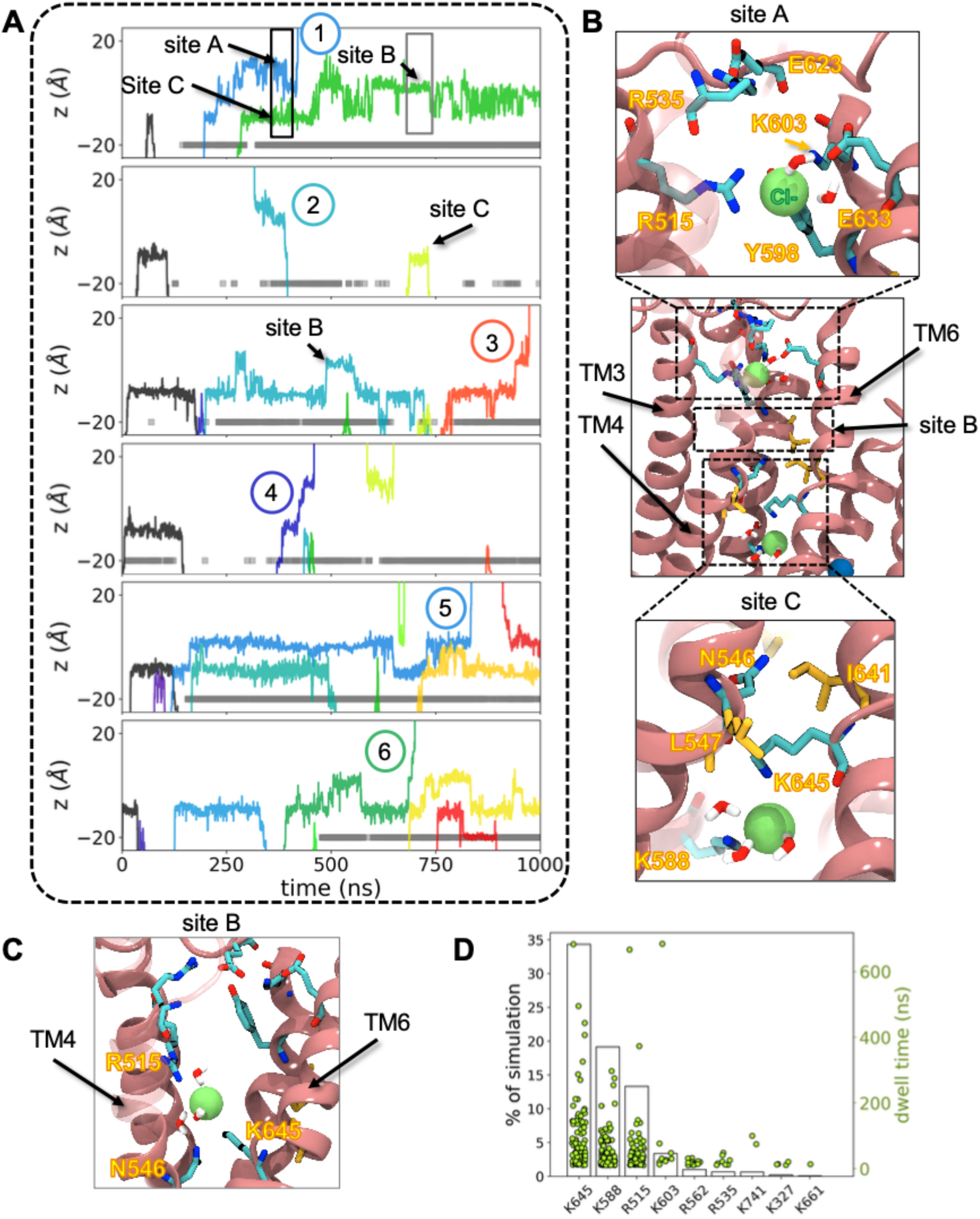
Spontaneous chloride permeation through dilated states of TMEM16A. **(A)** Z-positions of chloride ions in the pore, each color represents a unique ion. The y-axis is zeroed at the cryo-EM z-position of the L547 Cα. Only simulations containing complete permeation events (numbered) are shown. Event 2 is the only inward permeation. The gray bars indicate when the L547-I641 heavy atom distance is > 9 Å. **(B)** Snapshots of the open TMEM16A groove with zoomed-in images of two Cl^-^ interaction sites: site A (top) and site C (bottom). **(C)** Zoomed-in snapshot of a third Cl^-^ interaction site (site B). **(D)** Plot of the percentage of simulation time basic residues interact with Cl^-^ ions (left y-axis, black bars) and dwell times of each interaction instance (right y-axis, green dots).

From visual inspection of each permeation event, we see that the ion is typically accompanied by at least 3 water molecules, and it also directly interacts with specific positively charged residues lining the deepest part of the pore (Fig. 2B-C). K645, K588, and R515 spend 10-34% of the aggregate simulation time in contact with at least one Cl^-^ ion (Fig. 2D). Although most contact events are under 200 ns, interaction times with basic residues such as K645 (site B/C), K588 (site C), R515 (site A/B), and K603 (site A) can last longer. In contrast, residues near the pore entrances, such as R535 (above site A) and R562 (below site C), contact ions less frequently and for shorter times (Fig. 2D).

### Pore dilation maintains contact between TM4 and TM6 but breaks contact between TM3 and TM4

Our simulations yielded numerous changes in the protein upon 1PBC removal, and we applied dimensionality reduction using time-independent component analysis (tICA)^31^ to reveal the dominant conformations. Twenty Cα-Cα distance pairs across the pore domain (Fig. S8A) were used to analyze the entire data set treating each dimer subunit independently.

Subsequent Markov state modeling (MSM) on the output was then used to cluster configurations into 15 macrostates (Fig. S9 and Methods for details). Projecting each configuration into the first two tICs and analyzing the conformations of the cluster medoids reveals that the dominant motion (tIC 1, x-axis) is TM4-TM6 separation to open the pore, followed by the breaking of a network of hydrophobic residues in the TM3-TM4 loop (Fig. 3A, C). The second mode (tIC 2, y-axis) involves separation of the intracellular ends of TM3 and TM4. The points are color coded by density from high (yellow) to low (dark blue), and most conformations lie close to the 7ZK3 starting structure, with cluster 7 being the largest distinct configuration to its left along tIC 1. The medoids reveal key structural features already discussed such as the upper (yellow) and lower (cyan) TM4 kinks as well as new features like TM3-TM4 separation and tilting into the membrane (cluster 14) (Figs. S9 and S10).

**Figure 3.**
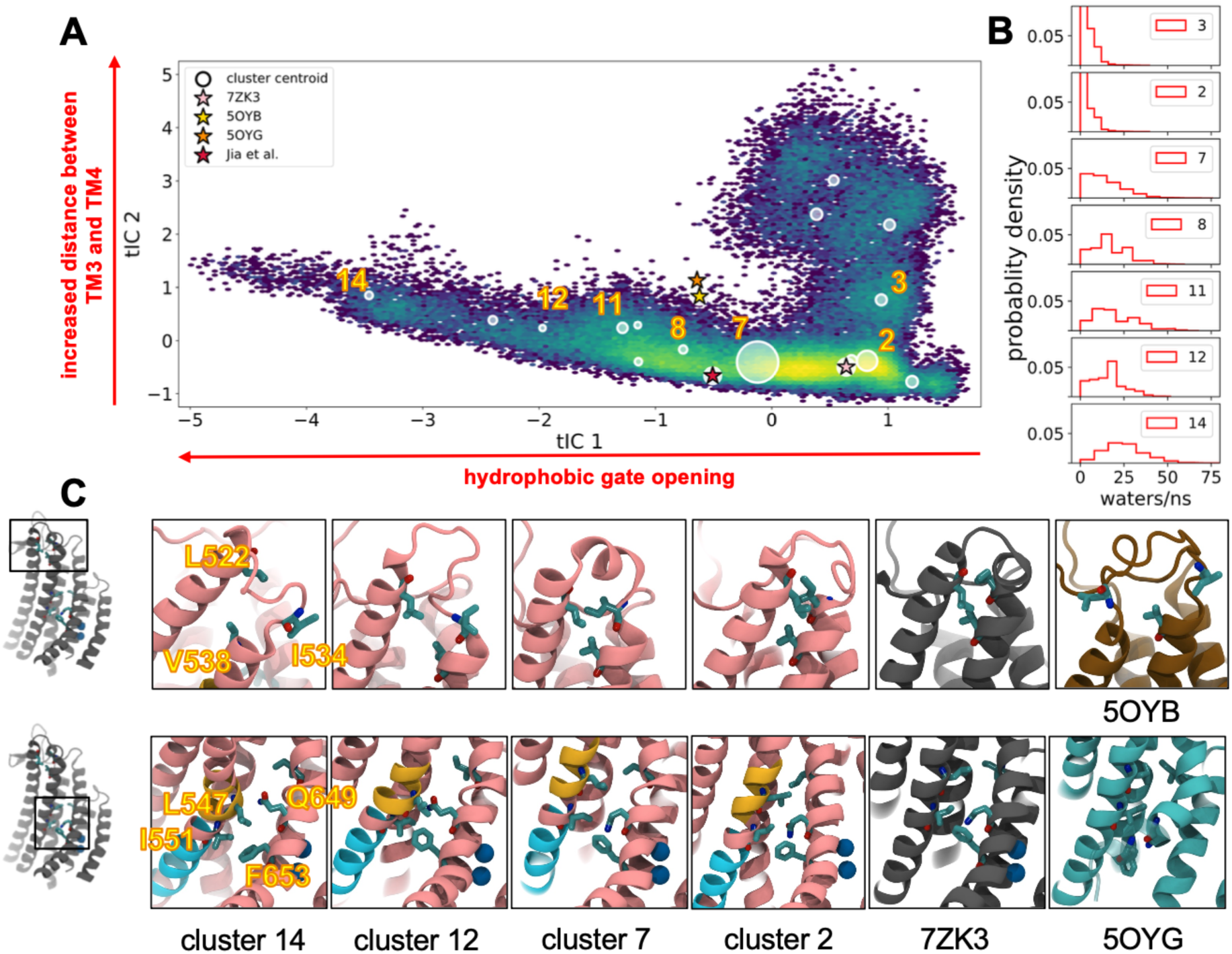
TMEM16A pore opening involved rearrangement of hydrophobic contacts. **(A)** All simulation data projected onto the first two tIC eigenvectors colored by log density. tIC 1 describes hydrophobic gate opening and tIC 2 distance between TM3 and TM4. Circles indicate cluster centroids sized by relative population; star denotes starting conformation. See Methods for details. **(B)** Histograms of water flux (number of waters/ns) through the center of the pore in select clusters. **(C)** Representative structures from selected clusters as labeled below each column. ***Top***: hydrophobic network at TM3/4 extracellular termini. ***Bottom***: hydrophobic contacts between TM4 and 6. Cryo-EM structures of 1PBC/Ca^2+^ -bound, Ca^2+^ -bound and apo TMEM16A shown in gray, brown, and cyan respectively. Structures for clusters 2, 7, and 14 are medoids. The non-pink colored portions of helix indicate locations on TM4 that bend to support opening (yellow) or further constricting (cyan) the channel.

We determined how open the pore is in each macrostate by computing the flux of water through snapshots in each cluster (Fig. 3B). Moving from right to left along tIC 1, the water flux increases from distributions peaked at 0 (cluster 2 and 3) to 25 events per ns (cluster 14) with the largest cluster (cluster 7) empirically marking the transition from a closed to open water pathway (Fig. 3B). We noticed that 50-70% of the subunits in the most hydrated clusters, 12 and 14, lost the α-helical structure at the extracellular end of TM4 and dissolved a small network of hydrophobic residues (L522 on TM3 and I534/V538 on TM4) (top row of Fig. 3C). In fact, cluster 14 additionally loses the upper kink at L547 and the extracellular end of TM4 separates from TM6 and moves into the membrane (yellow residues in Fig. S9). Thus, the hydrophobic network appears to anchor TM4 to TM3 to support the formation of the upper kink, and this network is stable as it remained intact in 98% of the aggregate simulation data. Finally, despite the extremely wide pores of clusters 12 and 14, TM4 and TM6 remain in close contact near the center of the pore through the formation of another hydrophobic interaction between L547 (TM4) and F653 (TM6) (Fig. S10, Fig. 3C, lower panel), which is discussed further below.

### A hydrophobic network in TM3 and TM4 stabilizes the active state

To probe the functional consequences of hydrophobic interactions at the external ends of TM3 and TM4 observed in our clustering analysis, we generated alanine mutants of L522 and I534 in TMEM16A and performed patch clamp electrophysiology on those constructs in HEK293 cells. Given that this network structure primarily occurred in medoids also demonstrating a pore sufficiently open to accommodate Cl^-^ ions, we reasoned that disrupting it by alanine substitution would disfavor the open state and lead to a shift in the voltage-dependence of activation. When intracellular pipette recording solutions included 600 nM Ca^2+^, we observed an outwardly rectifying pattern of current activation in wild-type TMEM16A when pulsed to a series of depolarizing voltages from a holding potential of -80 mV, which closed upon repolarization (Fig. 4A-B). When the same recording protocol was performed in TMEM16A L522A, the channel retained an outwardly rectifying pattern but showed markedly smaller currents which required stronger depolarization to exhibit measurable outward currents. To quantify this change, we normalized current values against the driving force for Cl^-^ ion flux (E_Cl-_ was 0 mV) to obtain macroscopic conductance, which we normalized to +150 mV before fitting G(V) curves using a two-state Boltzmann relationship. Mutation at L522A resulted in a rightward shift of the half-maximal activation (V_1/2_) of the G(V), from 42.3 ± 9.9 mV (n=9) to 137.7 ± 16.0 mV (n=5), consistent with a destabilization of the conducting confirmation by the mutant (p<.0001). A second mutation, I534A, exhibited an even more potent reduction in apparent current amplitude, with negligible outward current visible in 600 nM Ca^2+^ even at pulses to +150 mV (Fig. 4A). To facilitate opening in this construct, we recorded similar traces after increasing [Ca^2+^]_i_ to 2 μM. Under that condition, we were able to recover an outwardly rectifying phenotype (Fig. 4B), but normalized G(V) relationships still demonstrated a large hyperpolarizing shift in V_1/2_ to 113.7 ± 9.3 mV (n=7) compared with wild-type at the lower Ca^2+^ concentration (p<.001). Taken together, these results indicate that disruption of the TM3-4 hydrophobic network observed in larger diameter pore clusters results in instability of the open conformation of TMEM16A necessitating larger depolarizations or higher internal Ca^2+^ to achieve channel activation.

**Figure 4.**
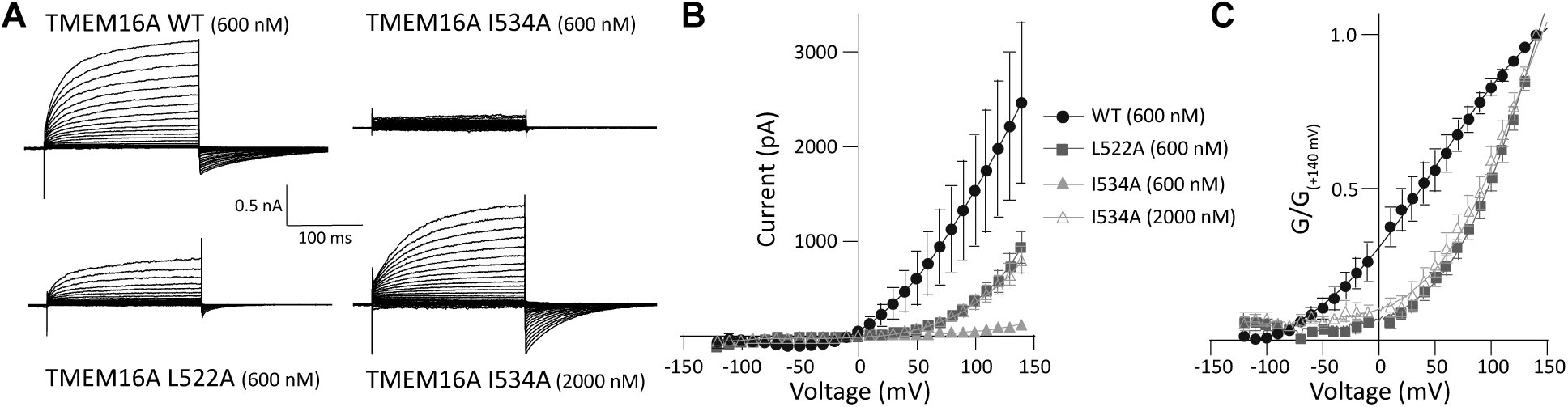
Mutations in the TM3-TM4 hydrophobic network shift TMEM16A activation gating. **(A)** Whole cell voltage clamp recordings from wild-type TMEM16A and L522A and I534A mutants expressed in HEK293 cells. Cells were held at -80 mV, then pulsed for 200 ms to potentials from -120 to +140 mV. As I534A exhibited negligible current amplitude at our initial base [Ca^2+^], a second, higher concentration was tested. **(B)** Current-voltage (I-V) plots for steady-state current amplitudes as in panel A. **(C)** Conductance-voltage (G-V) plots generated from data in A and normalized to the current amplitude at +140 mV. Two-state Boltzmann equations were fit to each trace. Conductance from the I534A mutant at the lower [Ca^2+^] was too low to properly fit a Boltzmann curve, so these data are not shown in panel C. n=4-8 for all conditions.

### Simulated current-voltage curve consistent with an open channel

The most dilated states of TM4 spontaneously conduct ions, but how these conformations correspond with known electrophysiological properties of the channel is unclear. We speculated that the diverse range of pore conformations seen in our clustering analysis may exhibit different simulated conduction properties, such as single channel conductance values, outward rectification, and so on. To answer this question, we performed MD simulations of representative conformations from 5 clusters in Fig. 3 under applied voltages ranging from -350 mV to +350 mV. Structures were taken from clusters 2, 8, 7, 11, and 12, and additional 1 μs long simulations at each voltage were initiated with backbone restraints to preserve the protein conformation (Fig. 5A and Table S2). These clusters were either highly represented in the aggregate data (2 and 7 are the largest) or sampled by simulations starting from different loop models (8, 11, and 12). They also exhibit key conformational features discussed already such as disengagement of the TM3-4 hydrophobic network (cluster 12) and bending at the upper TM4 kink (cluster 8).

**Figure 5.**
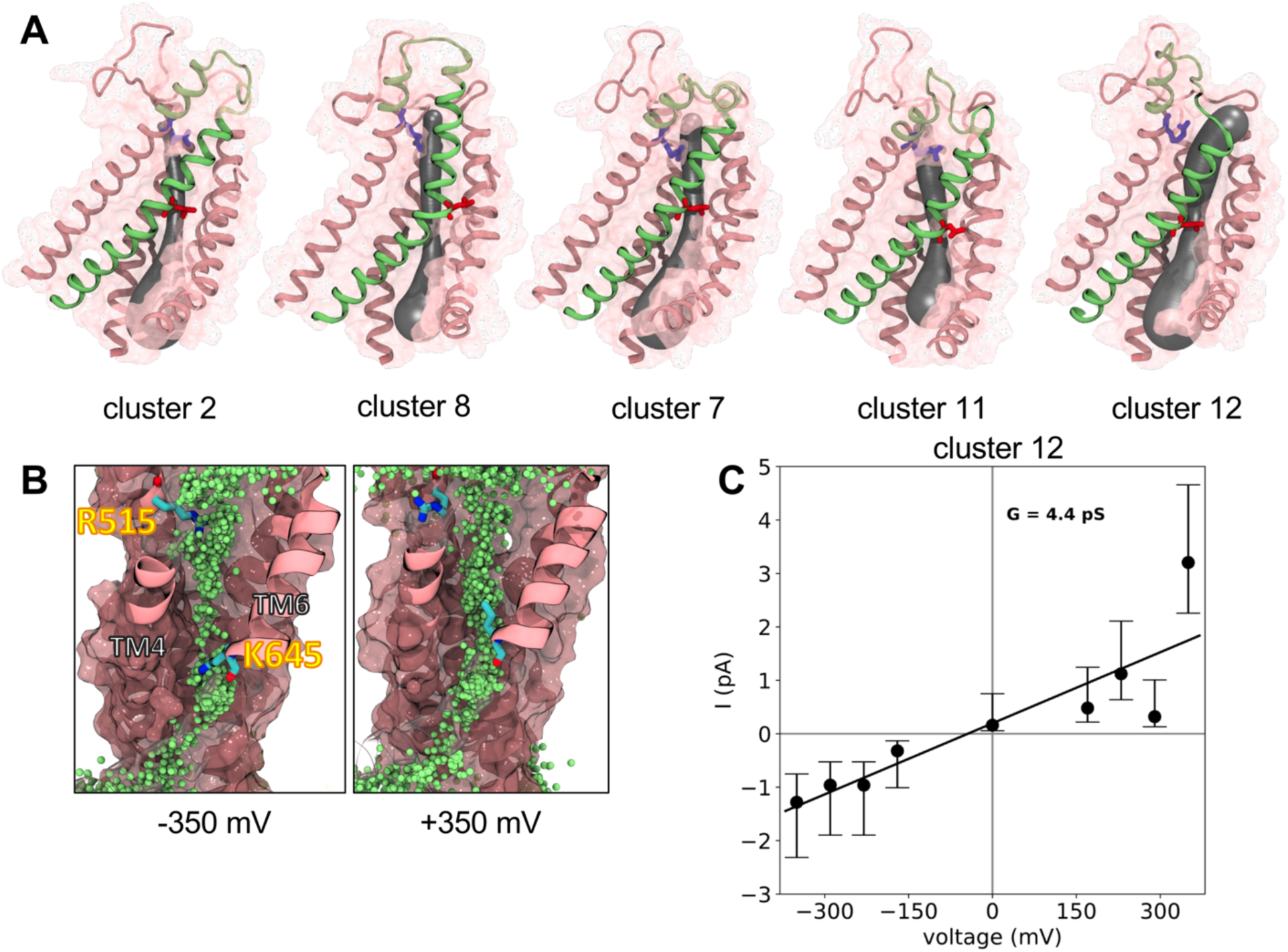
Permeation properties of several TMEM16A conformations. **(A)** Structures from TMEM16A clusters (MSM states): 2, 7, 8, 11 and 12. Only TM 3-8 are shown and TM4 is colored green without its solvent accessible surface shown (transparent pink elsewhere) and L547 (TM4, red) and R515 (TM3, blue) sidechains shown as sticks. Each structure shown with a 3D spline (gray) fit to the averaged pore center coordinates and radii from multiple simulation frames using HOLE2^34^. **(B)** Cartoon representation of the cluster 12 structure during simulations under an applied -350 mV (left) and 350 mV voltage with overlay of all Cl^-^ positions (in green) every 0.5 ns. **(C)** Currents calculated from simulations of cluster 12 plotted as a function of voltage. A linear fit was made to the current data and error bars indicate the 95% confidence interval taken from a Poisson distribution of event rates. Positional restraints were applied to the pore-lining helices of each simulation.

At 0 mV, simulations of the medoids from clusters 11, 8, 7, and 12 show a progressive increase in the *average* minimum pore radius from 1.1 to 2.0 Å, respectively, compared to 0.7 Å for cluster 2 close to the starting 1BPC-removed pore size (Fig. S12). Considering that the bare Cl^-^ ion radius is 1.8 Å, it is not surprising that only a single permeation event was captured across all of these conformations, namely in the one with the widest pore (cluster 12). In fact, across the entire voltage range clusters 2 and 8 showed no permeation events and cluster 7 showed only 2 events requiring a high positive voltage. We observed Cl^-^ throughout the pore at the most extreme voltages for clusters 12, 11, and to a lesser extent, 7, but cluster 8 showed little penetration (Fig. 5B and Fig. S13). In clusters 11 and 12, but not the others, K645 on TM6 adopted two rotamers depending on the voltage, but this does not appear to have an effect the stability of site B (Fig. S14, Videos S1-2).

Clusters 11 and 12 revealed 10 and 54 Cl^-^ permeation events, respectively, while K^+^ did not permeate any of the structures. At half of the voltages in the range, we were unable to simulate ion flow for cluster 11, but we saw events at each voltage in the low pA range for cluster 12 and used them to construct an I-V curve. A linear fit to the data corresponds to a 4.4 pS conductance (Fig. 5C) consistent with reports that range from 0.5 to 8 pS^2,5,32,33^.

Unfortunately, due to difficulties in simulating low conductance channels over the practical duration of MD times, ion counts at any specific voltage were low, leading to high uncertainty, and requiring us to use non-physiological voltages. Moreover, we observed highly asymmetric permeation counts at the most extreme voltages of ± 350 mV. Altogether, these factors made it difficult to determine if the channel is exhibiting Ohmic or outwardly rectifying behavior, though the presence of current at negative voltages in cluster 12 is more consistent with a linear (Ohmic) state than with the strong outward rectification behavior in moderate Ca^2+^. However, it remained possible that Ca^2+^ unbinding at negative voltages was controlling rectification.

### A kinetic model to explore the role of Ca^2+^ binding on conduction

To assess the influence of voltage and Ca^2+^ on the conduction behavior of the cluster 12 medoid, we computed the Cl^-^free energy profile along the pore (Fig. 6A-B) from the restrained MD simulations in the absence of a membrane voltage (black curve in Fig. 6B). Site C was the most energetically favorable, and was stabilized by about -2.5 kT with respect to the 150 mM Cl^-^ solution, while sites A and B were of equal energy with each other and solution. The largest barriers were the entrance to site C from the cytoplasm (∼3.5 kT barrier height) and between sites B and C (∼6 kT barrier moving from C to B), while A and B were separated by a barrier of about 2 thermal units. Profile asymmetry as seen in this energy profile is a feature that supports rectification, so we constructed a state-based kinetic model to explore the conduction properties of such a landscape. Assuming that at most 2 ions occupy the pore at a time and using rate constants constrained by the energetics in panel B, we devised a 3-site model with 7 states (panel C) with the full set of equations presented in Supplemental Kinetic Model with parameters in Tables S3-S4. Solving this system of equations with the energy profile corresponding to 2 bound Ca^2+^, we observed an Ohmic single-channel, current-voltage curve (black curve in Fig. 6D) as suggested by the MD simulations (Fig. 5C).

**Figure 6.**
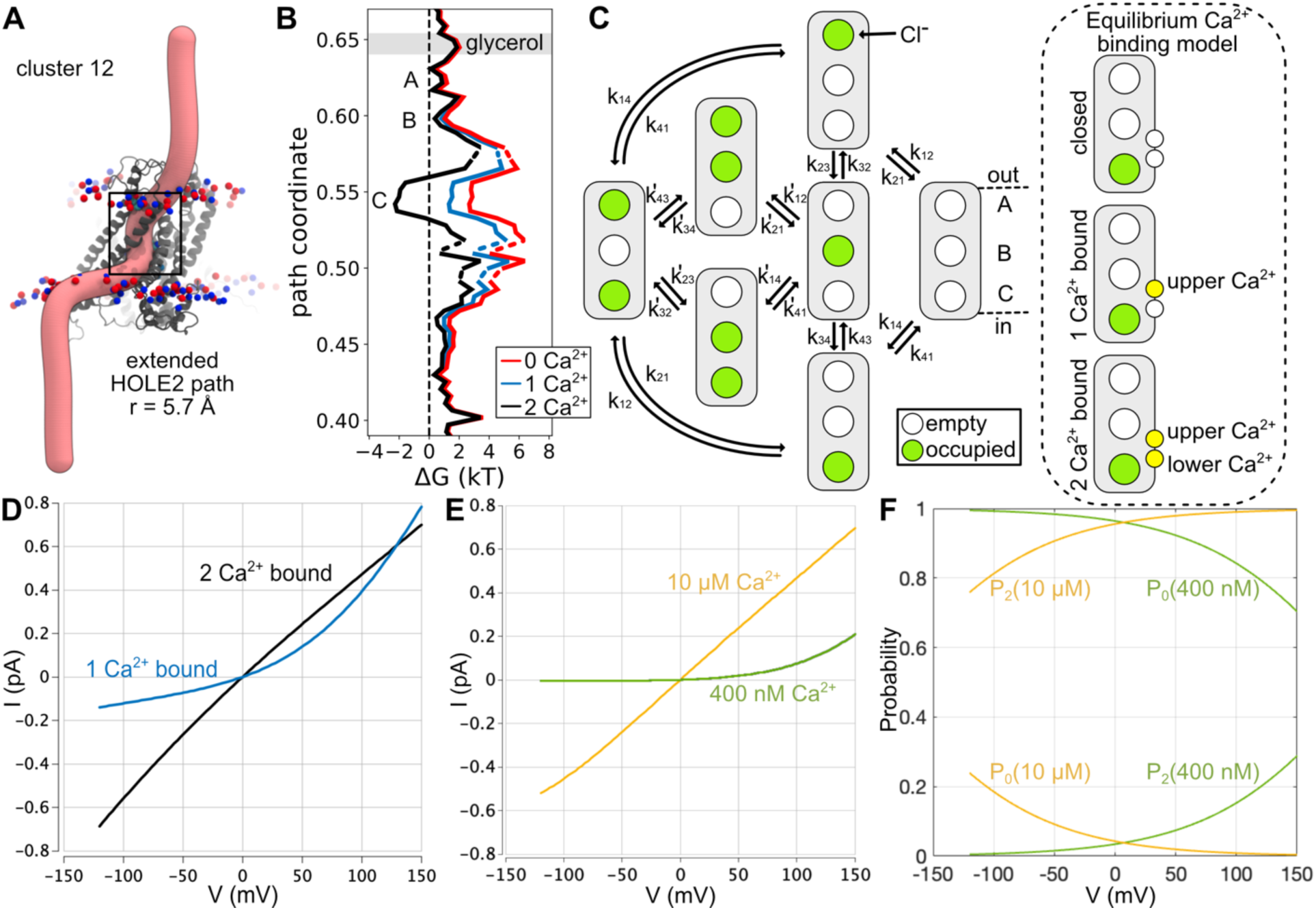
Cl^-^ permeation energies are strongly influenced by Ca^2+^ binding and can switch from Ohmic to rectifying conduction states. **(A)** The permeation pathway (pink tube) through the cluster 12 representative structure (gray) predicted with HOLE2 fit to a 2D spline and extended manually into solution. Phosphorous (red) and nitrogen (blue) atoms of the POPC headgroups are shown. **(B)** Free energy profiles for Cl^-^ permeation calculated from ion density along the pathway in A with both Ca^2+^ ions bound (black), the electrostatic component of the lower Ca^2+^ removed (blue), or the electrostatic components of both Ca^2+^ removed (red). **(C)** A 3-site kinetic model for Cl^-^ permeation based on observations from our simulations. Currents are estimates of single-channel values in 140 mM symmetric Cl^-^. Only 2 Cl^-^ (green circles) occupy the pore at any given time and the rate constants obey detailed balance based on the energy profiles in panel B. The Cl^-^ permeation model was coupled to an equilibrium Ca^2+^ (yellow circles) binding model (dashed oval) to determine the probability of the channel to be in the 0, 1, or 2 Ca^2+^ bound state. Supplemental Kinetic Model for details and all parameters. **(D)** Single channel currents from model with 2 Ca^2+^ bound (black) or 1 Ca^2+^ bound (blue). **(E)** Predicted ensemble averaged, single channel currents from model in panel C along with the equilibrium Ca^2+^ binding model. The model was computed at moderate 400 nM (green) and high 10 μM (yellow) intracellular Ca^2+^ levels. **(F)** The probability that the channel adopts the 0 Ca^2+^ (P_0_) and 2 Ca^2+^ (P_2_) bound states in intermediate (green curves) and high (yellow curves) calcium. The singly bound Ca^2+^ state is negligible in both conditions.

Previous studies have proposed that Ca^2+^ gates Cl^-^ conduction in constitutively active mutants electrostatically by neutralizing the negative charge in the Ca^2+^ binding site^35^. We tested this idea with our kinetic model of the wild-type channel by using continuum electrostatics calculations to predict the contribution of each Ca^2+^ to the Cl^-^ permeation profile in panel B (see Methods). Removing the lower Ca^2+^ (closer to the cytoplasm) destabilized site C and alters the barriers for getting into and out of the site (blue curve), consistent with calculations on the Ca^2+^ bound structure (5OYB)^35^. However, removing the upper Ca^2+^ as well (red curve) had little additional affect due to the non-linearity of the Poisson-Boltzmann equation, which is amplified by the high charge content of the intracellular pore and Ca^2+^ binding sites (Fig. S15). Hence, binding of the second Ca^2+^ has the greatest influence on the Cl^-^ permeation kinetics. We created a second kinetic model using the scheme in Fig. 6C with rate constants corresponding to the single Ca^2+^ bound profile (blue curve). The solution of this model is outwardly rectifying (blue curve in Fig. 6D), supporting the claim that electrostatics alone are a strong determinant of the permeation properties of TMEM16A^35^.

Nonetheless, two features of the I-V curves in panel D are inconsistent with the Ca^2+^ dependence of TMEM16A activation. First, switching from high to moderate Ca^2+^ causes a pronounced reduction in the total current at positive voltage^17^, but the 1 Ca^2+^ bound current is comparable to the 2 Ca^2+^ current at positive voltages and exceeds it above 125 mV. Second, the residual current at negative voltages is too large in the 1 Ca^2+^-bound state (compare to Fig. 4B). Our initial calculations ignored the voltage-dependence of Ca^2+^ binding, and we augmented the model (dashed oval in panel C) to include two sequential steps from a no-Ca^2+^-bound closed state to a single upper Ca^2+^-bound intermediate open state (blue curve in D) to a fully bound Ca^2+^ Ohmic state (black curve in D). The contribution of each state to the total cellular current is given by the fraction of channels in each state, as derived in the Supplemental Kinetic Model, and the fraction of the membrane electric field that the upper and lower Ca^2+^ experience upon binding was calculated with continuum electrostatics calculations. By tuning the otherwise unknown energetics of Ca^2+^ binding in the model, E_1_ (upper) and E_2_ (lower), with reasonable estimates and using the voltage dependence of binding estimated from electrostatic calculations (Table S5), the model reproduced these missing features as shown in Fig. 6E at 400 nM intermediate [Ca^2+^] (green curve) and saturating 10 μM [Ca^2+^] (yellow curve). In high calcium, the probability of 2 Ca^2+^ being bound (P_2_) is above 80% over the entire range and approaches 1 at depolarized values (yellow curves in Fig.6F). Meanwhile in 400 nM [Ca^2+^], the channel is closed at negative voltages with P_0_ ∼ 0, and only as voltage rises to 150 mV does P_2_ increase to ∼25% (lower green curve). This later curve resembles the I-V curves in sub-saturating conditions as in Fig. 4B.

### Lipids block ion permeation in most dilated state

Most other members of the TMEM16 family act as both lipid scramblases and ion channels and they share very similar structures to TMEM16A, including the location of the hydrophilic pathway for ions and lipids. By modeling ion-conductive states of TMEM16A we also gained an understanding of the structural details of how it acts solely as an ion channel. We did not observe any scrambling events during our simulations, but we did observe POPC headgroups penetrating deeply into the extracellular vestibule of the pore during simulations of cluster 12 (Fig. 7B). In other open states, the POPC headgroup would only rest at the edge of the extracellular pore with the choline facing and/or partially lining the ion pathway (Fig. 7E). In cases where the lipid headgroup entered the pore, we attributed a lack of scrambling to the close contact that is maintained between L547 on TM4 and F653 on TM6 (Fig. 3C), which in TMEM16 scramblases separate to allow lipids to pass^4^.

**Figure 7.**
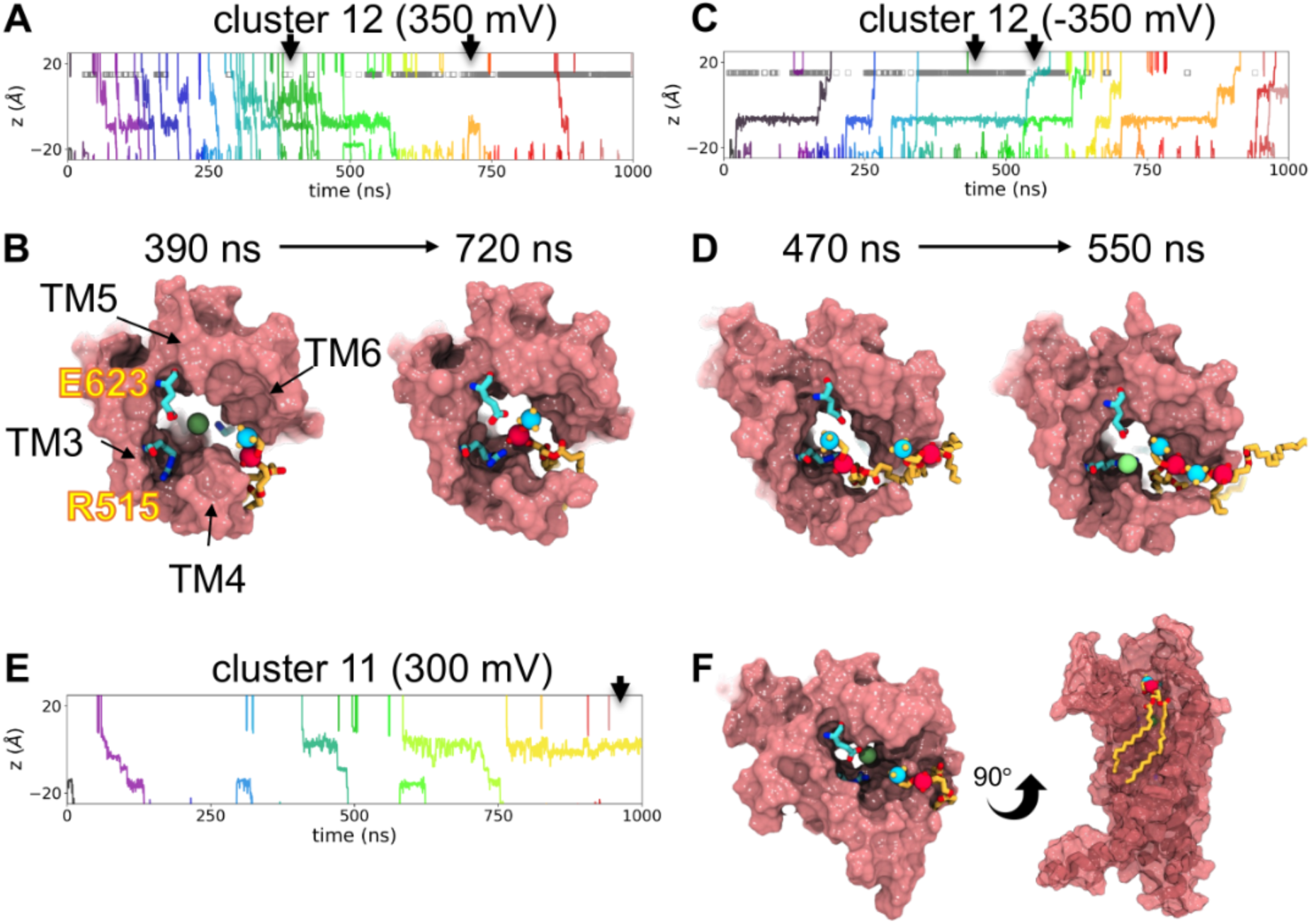
Lipids block ion permeation in most dilated TMEM16A states. **(A)** Traces of chloride z-positions in the TMEM16A pore during simulations of the cluster 12 structure with 350 mV applied voltage and **(B)** snapshots from the same simulation at timepoints indicated by the black arrows. **(C)** Traces of chloride z-positions in the TMEM16A pore during simulations of the cluster 12 structure with -350 mV applied voltage and **(D)** snapshots from the same simulation at timepoints indicated by the black arrows. **(E)** Traces of chloride z-positions in the TMEM16A pore during simulations of the cluster 12 structure with 350 mV applied voltage and **(F)** snapshots from the same simulation in panel E at timepoints indicated by the black arrow. The gray bars in panel A and C indicate when a lipid phosphorous atom is within 5 Å of the R515 sidechain. POPC lipids are shown in yellow, and chloride is shown as a green sphere.

Unexpectedly, when lipids do enter the pore, they block ion permeation in cluster 12 by interacting with the same basic residues that coordinate Cl^-^ in site A, namely R515 (Fig. 7A). This block is transient even on the short timescale of our simulations as occasionally the lipid fluctuates to allow ion passage (Fig. 7C). This opens new possibilities about how lipids are involved in ion conduction and inhibition even for non-scrambling TMEM16 members.

## Discussion

There are now at least 10 published structures of TMEM16A, seven of which are bound to Ca^2+^, however, none of them have a minimum pore radius large enough to accommodate a bare chloride ion (Table S1). Although the field has made strides to identify residues critical for conduction, we still do not know the open state of the pore and whether it adopts more than one conformation to support Ohmic and rectifying currents. One clue for what the Ohmic or fully conductive state might be comes from the 1PBC-bound structure, given that 1PBC is thought to stabilize the open channel state adopted in saturating calcium^26^. We hypothesized that removing 1PBC would allow the protein to relax to the fully conductive state in all-atom MD simulations. Across 12 unrestrained simulations, half of the 24 independent ion pores spontaneously dilated upon 1PBC removal, and subsequently we saw 6 spontaneous Cl^-^permeation events, which has not been reported in the absence of an applied voltage (Figs. 1-2).

Using tICA, clustering, and MSM analysis, we identified a number of conformational changes that occur within the protein as well as structural features that support ion passage once the pore dilates. First, permeation occurs once the gating residues L547 and I641 separate by more than 9 Å, and the upper portion of TM4 adopts a kink at residue L547 as recently reported by Kostritskii and co-workers, where they focused on the π-bulge that occurs to support the kink^30^. Cl^-^ ions reside at specific sites, termed A, B, and C, where basic residues stabilize the anion (Fig. 2B, C), and many of these pore-forming residues were predicted from mutagenesis^16,17,20^. Jia and Chen observed spontaneous pore opening and a single Cl^-^permeation event after placing PIP_2_ at an intracellular site composed of basic residues from TM3-5^23^. While one must take caution in estimating conductance values from a single event, or even the tens of events we observed at some voltages, they reported a value of 1.3 pS, which is slightly lower than the 4.4 pS we estimated for the I-V curve for cluster 12, but still in the 0.5-8.0 pS estimates from experiment^33^. They also reported a 6 kcal/mol Cl^-^ permeation barrier, while our calculations show a smaller barrier of 5.5 kT (3.3 kcal/mol) (Fig. 6B). We expect that our structure is wider, and while R515 was outside of the permeation pathway in their simulations, it is within the pathway in our open clusters and the structure presented in Kostritskii *et al.*^30^: this alone could explain their higher barrier. However, Jia and Chen also reported a higher anion selectivity than even the more selective outwardly rectifying condition, which we believe again indicates that their pore was not fully dilated.

The other 12 pores collapsed and dehydrated upon 1PBC removal. One explanation for these varied results is that 1PBC binds to and stabilizes a conformation along the activation gating trajectory that is intermediate between the fully closed and fully open states. In the absence of 1PBC this intermediate state may not be stable, allowing the protein to relax to a variety of new conformations. The most obvious structural feature in the transition toward a more closed pore is the formation of the lower kink on TM4 at E555 (Fig. S6). Additionally, it is possible that the pore of the 1PBC-bound structure may be in a “nearly open” conformation, but other factors, such as missing PIP_2_ or aspects of the intracellular domain, bias it toward a closed conformation. The importance of other protein elements in guiding the dynamics of the channel is highlighted in Fig. S11, where we show that the different intracellular loops influence movement through the tIC space. While all models sample clusters near the starting cryo-EM structure, including conductive pores like clusters 7 and 12, some regions of phase space are unique to particular model builds. Large values along tIC 2 (clusters 4-6) are only sampled by models without loops, and the most extreme motions along tIC 1 (clusters 11-14) are only sampled by models built using MODELLER. We speculate that these differences may be related to how the TM2-3 loop is modeled as it is adjacent to TM3 and 4 and can impact their dynamics, which as discussed, impacts gating. Thus, resolving these regions in structures and determining how they interact with PIP_2_ is important.

As cluster 12 exhibited spontaneous and bidirectional Cl^-^ permeation events, we used it as a representative open structure model to perform free energy and electrostatics calculations along the pore axis. We identified site C as the strongest electrostatic free energy well for Cl^-^, with asymmetric energy barriers above and below the site. It has been proposed that charge neutralization of the acidic residues at the Ca^2+^-binding sites upon Ca^2+^ binding accounts for the switch between rectifying and Ohmic currents^35^. By successively removing the Ca^2+^ ions from cluster 12, we calculated an increase in the electrostatic barriers flanking site C, and a marked shift in the energetic landscape at site C itself (3-5 kT), but minimal changes at sites A or B. The kinetic model with a single Ca^2+^ bound gives weak outward rectifying currents while the doubly bound Ca^2+^ model is Ohmic. Thus, our analysis of the model suggests that the rectifying current observed in experiment does not arise from channels with a single Ca^2+^ bound. Instead, rectification arises from the voltage-dependence of Ca^2+^ binding at low concentrations; only as the voltage becomes more positive are the ions driven into the binding site, opening the channels. These models support the notion that successive, voltage-dependent Ca^2+^-binding events shape the steady state current into rectifying or Ohmic forms, depending on the local [Ca^2+^].

Notably, the electrostatic profile of the pore did not vary appreciably between the 0 Ca^2+^ and 1 Ca^2+^-bound forms. This would imply that cluster 12 would conduct significant rectifying current even in the absence of Ca^2+^; however, we designated this conformation as non-conductive in our model to be consistent with experiment. The lack of Cl^-^ conductance in Ca^2+^-free channels likely stems from a gating transition from cluster 12 to a non-conductive state triggered by Ca^2+^ unbinding, perhaps to a state resembling the published apo conformation^21^ or our simulated medoids exhibiting smaller TM4-6 separations (Fig. S9). Moreover, the time scale of channel opening and closing at low activating [Ca^2+^] (e.g. Fig. 4A) is consistent with a voltage-dependent conformational change taking place at, or near, the Ca^2+^-binding site. For instance, in the absence of a coordinated Ca^2+^ ion, the acidic side chains of the binding site are likely to repel each other and may translate outward at negative voltages. The presence of a depolarizing voltage may draw the negatively charged site towards the inner leaflet into a more compatible conformation for Ca^2+^ binding, which could constitute a “primed” or “pre-open” state^17,26^. Transient association of the first Ca^2+^-ion may then serve to stabilize this confirmation and facilitate the positively cooperative second Ca^2+^ binding event. The primed state may also be what presents an outer pore binding site for 1PBC, which could then similarly stabilize the permeation pathway as observed in the 1PBC-bound structure^26^ and our subsequent simulations.

Consistent with this observation, our experiments identified a role for a small hydrophobic network at the extracellular termini of TM3 and 4 (Fig. 3), which is intact in the 1PBC-bound structure^26^ but eventually breaks as the protein samples the most dilated extracellular pore entrances (Fig. 3 and Fig. S12). Specifically, in clusters exhibiting an intact network, residues L522 and I534 were in direct contact. Conversely, L522 and I534 are oriented away from each other in structures with closed pores^21,22^. Reorientation within the TM3-4 linker and formation of this network may be concomitant with a gating motion from the closed state to the primed state. We propose that either 1PBC binding or membrane depolarization can enhance the primed state, and that transition into this confirmation precedes the first Ca^2+^ binding event. In line with this view, site directed mutagenesis of L522 or I534 resulted in a hyperpolarizing shift in the voltage-dependence of activation (Fig. 4). Moreover, TMEM16A channels with the stronger I534A mutation showed minimal activity at 600 nM Ca^2+^ even at +150 mV, and required 2 μM [Ca^2+^]_I_ to partially open, indicating higher [Ca^2+^]_I_ was needed to stabilize the primed state. The observations that elevating [Ca^2+^]_I_ enhances 1PBC potency^16,26^ and can also offset the destabilizing effects of I534A, suggest that Ca^2+^ binding and depolarization act in concert to favor states with an intact TM3-4 network. Additional experimental investigations of the process of voltage-dependent conformational transitions of TMEM16A in the absence of Ca^2+^ will further illuminate the complete order of events associated with channel gating.

MD simulations of TMEM16 scramblases demonstrate that lipid headgroups and ions share the same pathway and that lipids can scramble via the hydrophilic groove once TM4 and 6 have sufficiently separated^30,36–42^. Unlike open TMEM16 scramblase structures^37,43–45^ where TM4 separates from TM6 to dilate the path for lipid passage, in the TMEM16A simulations here, we see consistent close contact between TM4 and TM6 through formation of new contacts as the conformation of TM4 changes (Fig. 3C). We believe this property prevents scramblase activity in the conductive state of TMEM16A; however, we recently reported a single, rare scrambling event from a coarse-grained simulation of TMEM16A^36^, which may indicate that lipids can still scramble at very low undetectable rates, or it may be an inaccuracy in the coarse-grain force field. In the current set of all-atom MD simulations, we did not observe any lipid scrambling events despite deep headgroup penetration into the pore in simulations of the cluster 12 medoid; in fact lipids appear to temporarily block ion permeation (Fig. 7). We speculate that this block could be due to the smaller volume of the TMEM16A pore compared to the open groove of scramblases, or the potential higher affinity of sites on TMEM16A for lipid and ion interaction compared with equivalent positions along the scramblase grooves. So while it is plausible that lipids would also penetrate the TMEM16A pore, only fatty acids have been reported to inhibit TMEM16A and this mechanism of action is unknown^18^.

## METHODS

### TMEM16A simulation system preparation

All-atom simulations were initiated from the Ca^2+^/1PBC-bound mTMEM16A cryo-EM structure (PDB ID 7ZK3^26^) or an extracted snapshot from simulations of Ca^2+^ -bound mTMEM16A (PBD ID 5OYB) from Ref.^23^. Missing residues in 7ZK3 (260-266, 467-482, 526-527, and 669-682) were built using MODELLER v10.2^46^ or extracted from the mTMEM16A prediction deposited in the AlphaFold2 Protein Structure Database^28^. MODELLER-predicted loops were refined with LoopRefine^27^ using 10 iterations per residue. Loops extracted from the AlphaFold prediction where fit to the 7ZK3 after aligning on TM7-8. These loops were then annealed into the 7ZK3 structure by applying LoopRefine only to 3-4 resides bridging the inserted loops with the rest of the protein. PROPKA3^47^ was used to check the protonation state of protein residues. E624 and D405 are both weakly protonatable at neutral pH, but well solvated in the simulations, and therefore left in their negatively charged states. N and C termini were capped with methylamide and acetyl groups and then embedded in a 155×155 Å^2^ 1-palmitoyl-2-oleoyl-glycero-3-phosphocholine (POPC) bilayer and solvated in 150 mM KCl using CHARMM-GUI’s Membrane Builder^48^. System charges were neutralized using the same ions. Parameters for 1PBC were generated using the VMD Force Field Toolkit Plugin^49^ and Gaussian09^50^.

### MD simulations

Simulations were performed with Gromacs v2020.6^51^ with the CHARMM36^52^, CHARMM36m^53^, and TIP3P^54^ force fields for lipids, protein, and water, respectively. During minimization, equilibration and production distance restraints with 418.4 kJ×mol^-1^×nm^-2^ force constants were applied between the Cα atoms of residues 465 and 489, 454 and 566, 169 and 278, 126 and 176, 196 and 189, 123 and 282, 185 and 200 to stabilize the cytosolic domain.

Simulations were run using a 2 fs time step in an NPT ensemble. Temperature was set to 303.15 K using the Nosé-Hoover (45) thermostat with 1 ps piston decay constant. The semiisotropic pressure coupling was maintained at 1 bar using the Berendsen^55^ and Parrinello-Rahman^56^ barostats with a 5 ps piston decay constant and 4.5×10^-5^ compressibility during equilibration and production, respectively. Hydrogen atoms were constrained with LINCs^57^. Particle mesh Ewald (PME) was employed for long range electrostatics^58^. Van der Waals (VdW) interactions were smoothly switched to zero between 1.0 and 1.2 nm. A Verlet cut-off scheme was used for non-bonded interactions. The protein with its bound Ca^2+^ ions, the membrane, and solvent bath were treated as separate groups for the thermostat coupling and center of mass removal. Harmonic restraints of the protein backbone, sidechains, lipids and dihedrals were applied and slowly reduced over 8 equilibration steps totaling ∼32 ns. The equilibrated box size was ∼145×145×143 Å^3^. In simulations with restrained backbones a 418.4 kJ×mol^-1^×nm^-2^ harmonic restraint was applied to backbone heavy atoms of TM3 (residues 485-525), TM4 (residues 529-569), TM5 (residues 72-602), TM6 (residues 629-668) and TM7-8 (residues 699-745). Simulation coordinates were saved every 0.25 ns.

### tICA and clustering analysis

A set of 20 residue-to-residue minimum distances between TM3, TM4, TM5, and TM6 (shown in Fig. S9A, B) were extracted from the entire set of trajectories and submitted to time-independent component analysis (tICA)^31^ and subsequent MiniBatch K-means clustering into 500 clusters. There clusters were then used to construct a Markov State Model and subsequently grouped into 15 macrostates using the improved Perron-cluster cluster analysis (PCCA+) method^59,60^. The above analysis was performed using MSMBuilder v3.8^60^.

### MD simulations with an applied voltage

We embedded each structure extracted from clustering analysis in a new POPC bilayer and solvated following the same procedure described above in *TMEM16A simulation system preparation*. Voltage was increased in 10 mV jumps every 5 ns starting from the end of normal equilibration until the target voltage was reached. The potential was set using a constant electric field protocol where 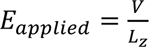 ^’^ where V is the voltage and L_z_ is the system box height at the end of normal equilibration (ranging between 152-161 Å). We simulated each structure for 1 μs each at -300, -250, -200, -150, 0, 150, 200, 250, and 300 mV (clusters 2, 7, 8, and 11) and -350, -290, -230, -170, 0, 170, 230, 290, and 350 mV (cluster 12 only). We applied 1 kcal/mol×Å^-2^ (kJ×mol^-1^×nm^-2^) restraints to the backbone heavy atoms of the pore-delineating TM3-8 backbone heavy atoms to maintain the starting conformation. The representative structure for cluster 12 was manually selected from one of the original simulations of the asymmetric model during a conduction event and later identified as a member of cluster 12. Other structures were either medoids (clusters 2, 7-8) or closest structure to the medoid of each cluster with modeled loops (cluster 11).

### Water and ion density calculations and pore pathway generation

The water and ion densities in the pore were calculated in a 25×35×150 Å^3^ rectangle centered on the Cα position of L547. Data from both subunits were aggregated after superposition, and the maps recorded at a 0.5 Å grid length resolution. The central pathway through the pore was then determined by calculating the weighted mean water density in slices along the z-axis through which a 3D spline was fit and nodes placed every 0.5 Å (ion density profiles in Fig. 1F,G) or 2.5 Å (water flux calculations in Fig. 3B) (see Fig. S3B).The ion density profiles in Fig. 1F,G were calculated along this water-defined pathway by counting ions within a 7 Å radial cutoff in z to the path (similar to the tube in Fig. 6A).

### Channel current and conductance measurements

The current for 200 ns blocks of simulation time was calculated as 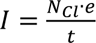 where N_Cl_ is the number of Cl^-^ permeation events, in the time block t, and e is the fundamental charge unit. The conductance in each block was calculated by dividing the current by the membrane potential. Mean and standard deviation of currents from each window were reported as the mean and error in Fig. 5C.

### Contact, flux, dwell time, and protein feature analysis

For every frame, the positions of all water and ions within a 12 Å radius of the average water density pathway were binned every 3.5 Å along the path. Interaction lifetimes were also recorded for any protein residue that came within 3.5 Å of an ion within the pore cutoff distance. A water flux event was counted every time a water molecule crossed the boundary between the cylindrical bin closest to the cryo-EM position of the L547 Cα into the adjacent bins. The contact analysis, water/ion position tracking and measurements for TM kink angles and residue distances were done using custom python scripts utilizing MDAnalysis v2.8^61^ and SciPy v1.14.1^62^ methods. Pore radii were calculated every 12.5 ns along the global z axis with sampling every 0.2 Å using the HOLE2^34^ implementation in MDAnalysis.

### Free energy and electrostatic potential calculations

Coordinates of the ion permeation pathway were calculated from the cluster 12 medoid in PDB format using HOLE2^34^. The base Cl^-^ free energy profile (black curve in Fig. 6B) was determined by taking the negative natural log of the ion density profile (as in Fig. 1D but with a small 5.7 Å radial cutoff) and setting the bulk energy to zero. To compute the electrostatic influence of bound Ca^2+^ ions on the permeating chloride anions and the influence of the membrane potential on the system, this same PDB structure was first converted to PQR format using PDB2PQR^63^ with the SWANSON^64^ partial charge parameter set. Zero, one, or two Ca^2+^ ions per dimer were reinserted into the PQR structure by hand at the known calcium binding sites. The electrostatic calculations were performed with APBSmem^65^ using parameters in Table 1. Modifications were made to the way APBSmem inserts the membrane into APBS^66^ input maps to ensure that the pathway was fully water solvated. Specifically, the membrane floodfill method used by APBSmem prevents infill of normally solvated cavities through small membrane-facing fenestrations by requiring the fenestration to be above a cutoff size, similar to a water probe radius but for lipids. This size was set to 15 Å. Due to high water content in the headgroup region observed in the MD, we assumed this region had a dielectric equal to water, and hence, we set the membrane thickness equal to that of the low-dielectric, hydrophobic core of the membrane. Additionally, the ion accessibility map and dielectric maps were modified along the identified pathway and the inner vestibule to ensure the pathway remained solvated using the results from the pathway identified with HOLE2. The SWANSON parameter set along with the 7th order surface spline option (srfm spl4) was chosen because it allows for good solution convergence near dielectric boundaries seen in a narrow pore. To calculate the membrane electrostatic field, we set all protein partial charges to zero and removed all ions and water from the system. We then applied membrane potential boundary conditions with 0 mV outside and -25 mV in the cytoplasm (u = -1 = eV/kT in reduced potential units at room temperature) to determine the electrostatic potential across the entire protein following gating charge calculations in APBSmem as described previously^67^.

**Table 1.** Parameters used for electrostatic potential calculation with APBSmem.

| parameter | value |
| --- | --- |
| partial charges | SWANSON |
| solution method | npbe or lpbe (as stated in the text) |
| boundary conditions | Zero or membrane potential |
| Grid dimensions | 161 x 161 x 161 |
| Grid length values | 300 $\text{\AA}$ x 300 $\text{\AA}$ x 300 $\text{\AA}$ |
| Focus lengths | 125 $\text{\AA}$ x 125 $\text{\AA}$ x 125 $\text{\AA}$ |
| Charge model | spl2 |
| Surface model | spl4 |
| srad | 1.4 $\text{\AA}$ |
| swin | 0.3 $\text{\AA}$ |
| sdens | 10 Å<sup>-2</sup> |
| temperature | 298.15 K |
| ions | symmetric 100 mM monovalent |
| Headgroup thickness | 0 Å |
| Membrane thickness (hydrophobic) | 29.2 Å |
| Solvent dielectric | 80 |
| Protein dielectric | 2 |
| Membrane insertion method | floodfill, with a 15 Å fenestration height limit |

Unlike our previous work, which treated the cytoplasmic effective charge density n_eff_ in simple geometric terms, we filled large cytoplasm-facing cavities, such as the hydrated vestibule up to the first constriction point, with the same the effective charge density n_eff_, making the entire cytoplasmic domain a conductor. Both non-linear and linear Poisson-Boltzmann calculations were performed to determine the potential along the pathway, but only the non-linear equation was employed in the membrane potential calculations and the energies in Fig. 6B employed non-linear form. In this case, n_eff_ (in Å^-3^) was set to n_eff_ = 2×10^-27^·N_A_·I·sinh(u), where N_A_ is Avogadro’s number, I is the molar concentration of the symmetric monovalent mobile salts, and u is the reduced membrane potential. The sinh term correctly propagates applied voltages of, i.e. -25 mV (-1 u), from the lower boundary up throughout the inner solvent space.

### cDNA material and cell culture

Wild-type mouse TMEM16A cDNA was a kind gift from Lily Y. Jan, University of California, San Francisco. TMEM16A cDNA was used as a fusion construct with eGFP, with the TMEM16A coding sequence cloned in-frame upstream of eGFP in the CMV-based mammalian expression plasmid pEGFP-N1. Point mutations were generated in TMEM16A by PCR using Phusion polymerase with 25-33 nucleotide custom mutagenic primers (IDT-DNA). Mutations were validated by Sanger sequencing (Genomics Research Core, UIC). HEK293T cells (ATCC) were seeded on T25 cell culture flasks and maintained in Dulbecco’s Modified Eagle’s Medium (DMEM) supplemented with 5% fetal bovine serum and 1% penicillin/streptomycin at 37°C and 5% CO_2_. HEK293 cells were routinely passaged with 0.25% Trypsin-EDTA. For experimental use, cells were seeded on 35 mm dishes for 24 hours prior to lipofection with Xtremegene 9 (Roche). 1-2 μg of cDNA was used to transfect each plate along with 4 μl of transfection reagent and allowed to incubate with cells for 24 hours. On the day of recording, cells were lifted for 3-5 minutes with Accutase (Gemini Bio), then plated in cell culture media onto 12 mm coverslips coated with 1% poly-L-lysine. Recordings were made 2-6 hours after plating. Cell culture reagents were from Gibco.

### Patch clamp electrophysiology

For patch clamp, cover slips seeded with HEK293 cells were transferred to a recording chamber containing a bath solution consisting of, in mM: 140 NaCl, 10 Hepes, 5 EGTA, adjusted to pH7.2 with NaOH and 308 mOsm with mannitol. Glass tips were pulled from filamented thick wall borosilicate glass (1.50/0.86, Sutter Instrument) using a P97 puller (Sutter) and polished to a tip resistance of 4-8 MΩ with a Narishige microforge. Pipette solutions contained, in mM: 140 NaCl, 10 Hepes, 2 MgCl_2_, and a mix of Ca_2_EGTA and EGTA calibrated to provide 600 nM or 2000 nM free Ca^2+^, as noted in the text, the adjusted to pH 7.2 with NaOH and 294 mOsm with mannitol. Free [Ca^2+^] in pipette solutions was measured directly by comparison to a fluorescent emission curve generated from known calcium standards, using Fluo-8 potassium salt dye (AAT Bioquest) and measured in a Molecular Devices SpectraMax plate reader. Whole cell patch clamp configuration was achieved by first obtaining a >GΩ seal, followed by manual aspiration until whole cell capacitance transients were observed. Whole cell capacitance was compensated, but no series resistance correction was used. Cells were held for >1 minute prior to recording to allow pipette solution to dialyze into the cell. Recordings were made with an Axoclamp 200B amplifier (Axon Instrument) and a Dendrite digitizer (Sutter) using a PC-based acquisition computer running SutterPatch3.0 software, and were sampled at 2 kHz. Conductance-voltage (G(V)) relationships were determined by dividing steady-state current amplitudes by the driving force for chloride flux at each applied voltage (V_m_ – E_Cl_), plotting these values against V_m_, and fitting each trace to a two-state Boltzmann function. As we could not identify one single [Ca^2+^]_I_ that was sufficient to elicit measurable current amplitudes for I534A that did not also lead to voltage-independent “Ohmic” current behavior in wild-type TMEM16A, G(V) curves were plotted at different [Ca^2+^]_i_ for these constructs: any shift likely underrepresents the impact of the mutation on voltage-dependence owing to the potentiating effect of the increased Ca^2+^ concentration used for I534A. Half-activation voltage (V_1/2_) for all traces for each condition were averaged and compared using one-way ANOVA followed by Tukey’s post-hoc test of multiple comparisons. Data are presented in the text as mean ± S.E.M., and p<0.05 was used as a threshold for statistical significance. Data analysis was performed offline with SutterPatch and Graphpad Prism software.

## Supporting information

Supplemental Text and Images

Zipped file of Mathematical Model (Berkeley Madonna format)

## ACKNOWLEDGEMENTS

We thank members of the Peters and Grabe laboratories for helpful discussion. This research was supported by the UCSF Discovery Fellows Program (C.A.S.), National Science Foundation Graduate Research Fellowship Program Grant No. 2038436 (C.A.S.), National Institutes of Health (NIH) R01 GM137109 (M.G.), R00 DA041500 (C.J.P.), and hardware for simulations was provided by grant no. NIH R01GM089740 (M.G.). Simulations were also carried out on the UCSF Wynton Cluster made possible through grant nos. NIH 1S10OD021596 and 1S10OD020054-011.

## AUTHOR CONTRIBUTIONS

C.A.S., C.J.P., and M.G. designed the project, C.A.S. performed all MD simulations and related analysis, C.J.P. performed all experiments, F.V.M performed the electrostatics calculations, M.G. performed the kinetic modeling, C.A.S. wrote the initial draft, and all authors wrote the manuscript. C.J.P. and M.G. supervised the project.

## REFERENCES

1. Schroeder, B.C., Cheng, T., Jan, Y.N., and Jan, L.Y. (2008). Expression Cloning of TMEM16A as a Calcium-Activated Chloride Channel Subunit. Cell 134, 1019–1029.

2. Yang, Y.D., Cho, H., Koo, J.Y., Tak, M.H., Cho, Y., Shim, W.S., Park, S.P., Lee, J., Lee, B., Kim, B.M., et al. (2008). TMEM16A confers receptor-activated calcium-dependent chloride conductance. Nature 455, 121–125.

3. Caputo, A., Caci, E., Ferrera, L., Pedemonte, N., Barsanti, C., Sondo, E., Pfeffer, U., Ravazzolo, R., Zegarra-Moran, O., and Galietta, L.J.V. (2008). TMEM16A, a membrane protein associated with calcium-dependent chloride channel activity. Science 322, 590–594.

4. Kalienkova, V., Mosina, V.C., and Paulino, C. (2021). The Groovy TMEM16 Family: Molecular Mechanisms of Lipid Scrambling and Ion Conduction. J. Mol. Biol. 433, 166941.

5. Zhu, M.H., Kim, T.W., Ro, S., Yan, W., Ward, S.M., Koh, S.D., and Sanders, K.M. (2009). A Ca^2+^-activated Cl^-^ conductance in interstitial cells of Cajal linked to slow wave currents and pacemaker activity. J. Physiol. 587, 4905–4918.

6. Jung, J., Nam, J.H., Park, H.W., Oh, U., Yoon, J.-H., and Lee, M.G. (2013). Dynamic modulation of ANO1/TMEM16A HCO₃^-^ permeability by Ca^2+^/calmodulin. Proc. Natl. Acad. Sci. U.S.A. 110, 360–365.

7. Cho, H., Yang, Y.D., Lee, J., Lee, B., Kim, T., Jang, Y., Back, S.K., Na, H.S., Harfe, B.D., and Wang, F. (2012). The calcium-activated chloride channel anoctamin 1 acts as a heat sensor in nociceptive neurons. Nat. Neurosci. 15, 1015–1021.

8. Huang, F., Zhang, H., Wu, M., Yang, H., Kudo, M., Peters, C.J., Woodruff, P.G., Solberg, O.D., Donne, M.L., Huang, X., et al. (2012). Calcium-activated chloride channel TMEM16A modulates mucin secretion and airway smooth muscle contraction. Proc. Natl. Acad. Sci. U.S.A. 109, 16354–16359.

9. Heinze, C., Seniuk, A., Sokolov, M.V., Huebner, A.K., Klementowicz, A.E., Szijártó, I.A., Schleifenbaum, J., Vitzthum, H., Gollasch, M., and Ehmke, H. (2014). Disruption of vascular Ca^2+^-activated chloride currents lowers blood pressure. J. Clin. Invest. 124, 675–686.

10. Chen, W., Gu, M., Gao, C., Chen, B., Yang, J., Xie, X., Wang, X., Sun, J., and Wang, J. (2021). The Prognostic Value and Mechanisms of TMEM16A in Human Cancer. Front. Mol. Biosci. 8, 542156.

11. Pinard, A., Ye, W., Fraser, S.M., Rosenfeld, J.A., Pichurin, P., Hickey, S.E., Guo, D., Cecchi, A.C., Boerio, M.L., and Guey, S. (2023). Rare variants in ANO1, encoding a calcium-activated chloride channel, predispose to moyamoya disease. Brain 146, 3616–3623.

12. Liang, P., Wan, Y.C.S., Yu, K., Hartzell, H.C., and Yang, H. (2024). Niclosamide potentiates TMEM16A and induces vasoconstriction. J. Gen. Physiol. 156, e202313460.

13. Tembo, M., Wozniak, K.L., Bainbridge, R.E., and Carlson, A.E. (2019). Phosphatidylinositol 4,5-bisphosphate (PIP2) and Ca^2+^ are both required to open the Cl^-^ channel TMEM16A. J. Biol. Chem. 294, 12556–12564.

14. Yu, K., Jiang, T., Cui, Y.Y., Tajkhorshid, E., and Hartzell, H.C. (2019). A network of phosphatidylinositol 4,5-bisphosphate binding sites regulates gating of the Ca^2+^-activated Cl^-^ channel ANO1 (TMEM16A). Proc. Natl. Acad. Sci. U.S.A. 116, 21050–21058.

15. Le, S.C., Jia, Z., Chen, J., and Yang, H. (2019). Molecular basis of PIP2-dependent regulation of the Ca^2+^-activated chloride channel TMEM16A. Nat. Commun. 10, 3729.

16. Peters, C.J., Yu, H., Tien, J., Jan, Y.N., Li, M., and Jan, L.Y. (2015). Four basic residues critical for the ion selectivity and pore blocker sensitivity of TMEM16A calcium-activated chloride channels. Proc. Natl. Acad. Sci. U.S.A. 112, 13826–13831.

17. Peters, C.J., Gilchrist, J.M., Tien, J., Bethel, N.P., Qi, L., Chen, T., Wang, L., Jan, Y.N., Grabe, M., and Jan, L.Y. (2018). The Sixth Transmembrane Segment Is a Major Gating Component of the TMEM16A Calcium-Activated Chloride Channel. Neuron 97, 1060–1073.

18. De Jesús-Pérez, J.J., Cruz-Rangel, S., Espino-Saldaña, Á.E., Martínez-Torres, A., Qu, Z., Hartzell, H.C., Corral-Fernandez, N.E., Pérez-Cornejo, P., and Arreola, J. (2018). Phosphatidylinositol 4,5-bisphosphate, cholesterol, and fatty acids modulate the calcium-activated chloride channel TMEM16A (ANO1). Biochim. Biophys. Acta Mol. Cell Biol. Lipids 1863, 299–312.

19. Leon-Aparicio, D., Sánchez-Solano, A., Arreola, J., and Perez-Cornejo, P. (2022). Oleic acid blocks the calcium-activated chloride channel TMEM16A/ANO1. Biochim. Biophys. Acta Mol. Cell Biol. Lipids 1867, 159134.

20. Paulino, C., Neldner, Y., Lam, A.K.M., Kalienkova, V., Brunner, J.D., Schenck, S., and Dutzler, R. (2017). Structural basis for anion conduction in the calcium-activated chloride channel TMEM16A. eLife 6, e26232.

21. Paulino, C., Kalienkova, V., Lam, A.K.M., Neldner, Y., and Dutzler, R. (2017). Activation mechanism of the calcium-activated chloride channel TMEM16A revealed by cryo-EM. Nature 552, 421–425.

22. Dang, S., Feng, S., Tien, J., Peters, C.J., Bulkley, D., Lolicato, M., Zhao, J., Zuberbühler, K., Ye, W., Qi, L., et al. (2017). Cryo-EM structures of the TMEM16A calcium-activated chloride channel. Nature 552, 426–429.

23. Jia, Z., and Chen, J. (2021). Specific PIP2 binding promotes calcium activation of TMEM16A chloride channels. Commun. Biol. 4, 1152.

24. Lam, A.K., and Dutzler, R. (2023). Mechanistic basis of ligand efficacy in the calcium-activated chloride channel TMEM16A. EMBO J. 42, e115030.

25. Lam, A.K.M., Rheinberger, J., Paulino, C., and Dutzler, R. (2021). Gating the pore of the calcium-activated chloride channel TMEM16A. Nat. Commun. 12, 785.

26. Lam, A.K.M., Rutz, S., and Dutzler, R. (2022). Inhibition mechanism of the chloride channel TMEM16A by the pore blocker 1PBC. Nat. Commun. 13, 1234.

27. Fiser, A., and Sali, A. (2003). ModLoop: Automated modeling of loops in protein structures. Bioinformatics 19, 250–251.

28. Jumper, J., Evans, R., Pritzel, A., Green, T., Figurnov, M., Ronneberger, O., Tunyasuvunakool, K., Bates, R., Žídek, A., Potapenko, A., et al. (2021). Highly accurate protein structure prediction with AlphaFold. Nature 596, 583–589.

29. Lam, A.K.M., and Dutzler, R. (2021). Mechanism of pore opening in the calcium-activated chloride channel TMEM16A. Nat. Commun. 12, 786.

30. Kostritskii, A.Y., Kostritskaia, Y., Dmitrieva, N., Stauber, T., and Machtens, J.-P. (2025). Calcium-activated chloride channel TMEM16A opens via pi-helical transition in transmembrane segment 4. Proc. Natl. Acad. Sci. U.S.A. 122, e2421900122.

31. Pérez-Hernández, G., Paul, F., Giorgino, T., Fabritiis, G.D., and Noé, F. (2013). Identification of slow molecular order parameters for Markov model construction. J. Chem. Phys. 139, 015102.

32. Takahashi, T., Neher, E., and Sakmann, B. (1987). Rat brain serotonin receptors in Xenopus oocytes are coupled by intracellular calcium to endogenous channels. Proc. Natl. Acad. Sci. U.S.A. 84, 5063–5067.

33. Whitlock, J.M., and Hartzell, H.C. (2016). A Pore Idea: the ion conduction pathway of TMEM16/ANO proteins is composed partly of lipid. Pflügers Arch. 468, 455–473.

34. Smart, O.S., Neduvelil, J.G., Wang, X., Wallace, B.A., and Sansom, M.S.P. (1996). HOLE: A program for the analysis of the pore dimensions of ion channel structural models. J. Mol. Graph. 14, 354–360.

35. Lam, A.K., and Dutzler, R. (2018). Calcium-dependent electrostatic control of anion access to the pore of the calcium-activated chloride channel TMEM16A. eLife 7, e39122.

36. Stephens, C.A., Hilten, N. van, Zheng, L., and Grabe, M. (2025). Simulation-based survey of TMEM16 family reveals that robust lipid scrambling requires an open groove. eLife 14, e105111.

37. Bushell, S.R., Pike, A.C.W., Falzone, M.E., Rorsman, N.J.G., Ta, C.M., Corey, R.A., Newport, T.D., Christianson, J.C., Scofano, L.F., Shintre, C.A., et al. (2019). The structural basis of lipid scrambling and inactivation in the endoplasmic reticulum scramblase TMEM16K. Nat. Commun. 10, 3956.

38. Stansfeld, P.J., Goose, J.E., Caffrey, M., Carpenter, E.P., Parker, J.L., Newstead, S., and Sansom, M.S.P. (2015). MemProtMD: Automated Insertion of Membrane Protein Structures into Explicit Lipid Membranes. Structure 23, 1350–1361.

39. Bethel, N.P., and Grabe, M. (2016). Atomistic insight into lipid translocation by a TMEM16 scramblase. Proc. Natl. Acad. Sci. U.S.A. 113, 14049–14054.

40. Lee, B.-C., Khelashvili, G., Falzone, M., Menon, A.K., Weinstein, H., and Accardi, A. (2018). Gating mechanism of the extracellular entry to the lipid pathway in a TMEM16 scramblase. Nat. Commun. 9, 3251.

41. Le, T., Jia, Z., Le, S.C., Zhang, Y., Chen, J., and Yang, H. (2019). An inner activation gate controls TMEM16F phospholipid scrambling. Nat. Commun. 10, 1051.

42. Khelashvili, G., Kots, E., Cheng, X., Levine, M.V., and Weinstein, H. (2022). The allosteric mechanism leading to an open-groove lipid conductive state of the TMEM16F scramblase. Commun. Biol. 5, 990.

43. Brunner, J.D., Lim, N.K., Schenck, S., Duerst, A., and Dutzler, R. (2014). X-ray structure of a calcium-activated TMEM16 lipid scramblase. Nature 516, 207–212.

44. Kalienkova, V., Mosina, V.C., Bryner, L., Oostergetel, G.T., Dutzler, R., and Paulino, C. (2019). Stepwise activation mechanism of the scramblase nhTMEM16 revealed by cryo-EM. eLife 8, e44364.

45. Falzone, M.E., Rheinberger, J., Lee, B.C., Peyear, T., Sasset, L., Raczkowski, A.M., Eng, E.T., Lorenzo, A.D., Andersen, O.S., Nimigean, C.M., et al. (2019). Structural basis of Ca2+-dependent activation and lipid transport by a TMEM16 scramblase. eLife 8, e43229.

46. Šali, A., and Blundell, T.L. (1993). Comparative protein modelling by satisfaction of spatial restraints. J. Mol. Biol.234, 779–815.

47. Olsson, M.H.M., Søndergaard, C.R., Rostkowski, M., and Jensen, J.H. (2011). PROPKA3: Consistent treatment of internal and surface residues in empirical pKa predictions. J. Chem. Theory Comput. 7, 525–537.

48. Jo, S., Lim, J.B., Klauda, J.B., and Im, W. (2009). CHARMM-GUI membrane builder for mixed bilayers and its application to yeast membranes. Biophys. J. 97, 50–58.

49. Humphrey, W., Dalke, A., and Schulten, K. (1996). VMD: Visual molecular dynamics. J. Mol. Graph. 14, 33–38.

50. Frisch, M.J., Trucks, G.W., Schlegel, H.B., Scuseria, G.E., Robb, M.A., Cheeseman, J.R., Scalmani, G., Barone, V., Mennucci, B., Petersson, G.A., et al. (2009). Gaussian∼09 Revision E.01.

51. Abraham, M.J., Murtola, T., Schulz, R., Páll, S., Smith, J.C., Hess, B., and Lindahl, E. (2015). GROMACS: High performance molecular simulations through multi-level parallelism from laptops to supercomputers. SoftwareX 1–2, 19–25.

52. Huang, J., and MacKerell, A.D. (2013). CHARMM36 all-atom additive protein force field: Validation based on comparison to NMR data. J. Comput. Chem. 34, 2135–2145.

53. Huang, J., Rauscher, S., Nawrocki, G., Ran, T., Feig, M., Groot, B.L.D., Grubmüller, H., and MacKerell, A.D. (2016). CHARMM36m: An improved force field for folded and intrinsically disordered proteins. Nat. Methods 14, 71–73.

54. Jorgensen, W.L., Chandrasekhar, J., Madura, J.D., Impey, R.W., and Klein, M.L. (1983). Comparison of simple potential functions for simulating liquid water. J. Chem. Phys. 79, 926–935.

55. Berendsen, H.J.C., Postma, J.P.M., van Gunsteren, W.F., DiNola, A., and Haak, J.R. (1984). Molecular dynamics with coupling to an external bath. J. Chem. Phys. 81, 3684–3690.

56. Parrinello, M., and Rahman, A. (1981). Polymorphic transitions in single crystals: A new molecular dynamics method. J. Appl. Phys. 52, 7182–7190.

57. Hess, B., Bekker, H., Berendsen, H.J.C., and Fraaije, J.G.E.M. (1997). LINCS: A linear constraint solver for molecular simulations. J. Comput. Chem. 18, 1463–1472.

58. Essmann, U., Perera, L., Berkowitz, M.L., Darden, T., Lee, H., and Pedersen, L.G. (1995). A smooth particle mesh Ewald method. J. Chem. Phys. 103, 8577–8593.

59. Deuflhard, P., and Weber, M. (2005). Robust Perron cluster analysis in conformation dynamics. Linear Algebra Appl.398, 161–184.

60. Harrigan, M.P., Sultan, M.M., Hernández, C.X., Husic, B.E., Eastman, P., Schwantes, C.R., Beauchamp, K.A., McGibbon, R.T., and Pande, V.S. (2017). MSMBuilder: Statistical models for biomolecular dynamics. Biophys. J. 112, 10–15.

61. Michaud-Agrawal, N., Denning, E.J., Woolf, T.B., and Beckstein, O. (2011). MDAnalysis: A toolkit for the analysis of molecular dynamics simulations. J. Comput. Chem. 32, 2319–2327.

62. Virtanen, P., Gommers, R., Oliphant, T.E., Haberland, M., Reddy, T., Cournapeau, D., Burovski, E., Peterson, P., Weckesser, W., Bright, J., et al. (2020). SciPy 1.0: fundamental algorithms for scientific computing in Python. Nat. Methods 17, 261–272.

63. Dolinsky, T.J., Nielsen, J.E., McCammon, J.A., and Baker, N.A. (2004). PDB2PQR: an automated pipeline for the setup of Poisson-Boltzmann electrostatics calculations. Nucleic Acids Res. 32, W665–W667.

64. Swanson, J.M.J., Wagoner, J.A., Baker, N.A., and McCammon, J.A. (2007). Optimizing the Poisson dielectric boundary with explicit solvent forces and energies: Lessons learned with atom-centered dielectric functions. J. Chem. Theory Comput. 3, 170–183.

65. Marcoline, F.V., Bethel, N., Guerriero, C.J., Brodsky, J.L., and Grabe, M. (2015). Membrane protein properties revealed through data-rich electrostatics calculations. Structure 23, 1523–1533.

66. Jurrus, E., Engel, D., Star, K., Monson, K., Brandi, J., Felberg, L.E., Brookes, D.H., Wilson, L., Chen, J., Liles, K., et al. (2018). Improvements to the APBS biomolecular solvation software suite. Protein Sci. 27, 112–128.

67. Callenberg, K.M., Choudhary, O.P., Forest, G.L., Gohara, D.W., Baker, N.A., and Grabe, M. (2010). APBSmem: A graphical interface for electrostatic calculations at the membrane. PLoS ONE 5, e12722.

