## Supplemental Text and Images for "Conformational changes upon pore blocker removal reveal conductive states of TMEM16A"

### Supplemental Kinetic Model

#### 1 A 3-site kinetic model of chloride permeation

$$
\begin{aligned}
\frac{d\,XXX}{dt} &= -(k_{12}C_{\text{out}} + k_{14}C_{\text{in}})\,XXX + k_{41}\,XXC + k_{21}\,AXX \\
\frac{d\,AXX}{dt} &= -(k_{21} + k_{23} + k_{14}C_{\text{in}})\,AXX + k_{12}C_{\text{out}}\,XXX + k_{32}\,XBX + k_{41}\,AXC \\
\frac{d\,XBX}{dt} &= -(k_{32} + k_{34} + k_{12}C_{\text{out}} + k_{14}C_{\text{in}})\,XBX + k_{23}\,AXX + k_{43}\,XXC + k_{21}\,ABX + k_{41}\,XBC \\
\frac{d\,XXC}{dt} &= -(k_{43} + k_{41} + k_{12}C_{\text{out}})\,XXC + k_{34}\,XBX + k_{14}C_{\text{in}}\,XXX + k_{21}\,AXC \\
\frac{d\,XBC}{dt} &= -(k_{41} + k_{32})\,XBC + k_{14}C_{\text{in}}\,XBX + k_{23}\,AXC \\
\frac{d\,ABX}{dt} &= -(k_{21} + k_{34})\,ABX + k_{12}C_{\text{out}}\,XBX + k_{43}\,AXC \\
\frac{d\,AXC}{dt} &= -(k_{21} + k_{41} + k_{23} + k_{43})\,AXC + k_{14}C_{\text{in}}\,AXX + k_{12}C_{\text{out}}\,XXC + k_{32}\,XBC + k_{34}\,ABX
\end{aligned}
$$

$XXX$, $AXX$, $XBX$, $XXC$, $ABX$, $AXC$ and $XBC$ are the occupancy probabilities of all possible states consisting of 0, 1, or 2 $Cl^-$ in the pore, where $A$, $B$ or $C$ as the first, second, and third character, respectively, indicates an ion is present at that site (see main text for definitions), while $X$ in those positions indicates an unoccupied site. The occupancy probabilities sum to one. $C_{\text{in/out}}$ are the $Cl^-$ concentrations in the cytoplasm (in) and extracellular space (out). The model is three site since ions can exist at one of three sites ($A$, $B$, or $C$), and we only allow transitions of a single ion at a time between adjacent sites. The transition rates ($k_{ij}$) have subscripts $i$ and $j$ representing movement of $Cl^-$ *from* site $i$ *to* site $j$, where site $A$, $B$, and $C$ correspond to subscripts 2, 3, 4, respectively, and the ion in solution corresponds to subscript 1. Rate constants $k_{12}$ and $k_{14}$ are second order and all other rates are first order. We then use Eyring rate theory to describe the dependence of the rates on the membrane voltage $V$:

$$
\begin{aligned}
k_{12} &= k_0' \exp\bigl(-E_{\text{out},A} - \varepsilon_{12}V_r\bigr) & k_{21} &= k_0 \exp\bigl(-(E_{\text{out},A} - E_A) - \varepsilon_{21}V_r\bigr) \\
k_{23} &= k_0 \exp\bigl(-(E_{A,B} - E_A) - \varepsilon_{23}V_r\bigr) & k_{32} &= k_0 \exp\bigl(-(E_{A,B} - E_B) - \varepsilon_{32}V_r\bigr) \\
k_{34} &= k_0 \exp\bigl(-(E_{B,C} - E_B) - \varepsilon_{34}V_r\bigr) & k_{43} &= k_0 \exp\bigl(-(E_{B,C} - E_C) - \varepsilon_{43}V_r\bigr) \\
k_{41} &= k_0 \exp\bigl(-(E_{\text{C,in}} - E_C) - \varepsilon_{41}V_r\bigr) & k_{14} &= k_0' \exp\bigl(-E_{\text{C,in}} - \varepsilon_{14}V_r\bigr)
\end{aligned}
$$

where $V_r = FV/RT$ is the reduced membrane potential at room temperature $T$, $F$ is the Faraday constant and $R$ is the gas constant, $(\varepsilon_{ij} - \varepsilon_{ji})$ is the equivalent charge movement between sites $i$ and $j$ based on our electrostatics calculations, we assume $\varepsilon_{ij} = -\varepsilon_{ji}$, $E_i$ is the $Cl^-$ energy at site $i$, $E_{i,j}$ is the barrier energy for moving from site $i$ to $j$, and $k_0$ and $k_0'$ are prefactors for ion movement. The rates $k_0'$ and $k_0$ have units that give solutions in ions per microsecond, and their values were adjusted to best match the unitary currents from our MD simulations and estimates from the literature (see Table S4). The quantity $2\varepsilon_{i,j}$ is the fraction of the membrane electric field experienced by moving from state $i$ to $j$, and $2(\varepsilon_{14} + \varepsilon_{43} + \varepsilon_{32} + \varepsilon_{21}) = +1$ reflects that a single ion moving through the entire channel experiences the full membrane voltage.

The rate constants obey detailed balance based on the energetics, and the energy of the anion in each state (in $kT$) was determined from the $Cl^-$ permeation profiles in Fig. 6B:

$$E_A = 0$$
$$E_B = 0$$
$$E_C = 3 - 1.5\,\mathrm{Ca}_1 - 3.5\,\mathrm{Ca}_2$$

where $\mathrm{Ca}_1$ and $\mathrm{Ca}_2$ are zero or one depending on if the lower or upper $Ca^{2+}$ are bound, respectively. Hence, when both calcium ions are present $\mathrm{Ca}_1$ and $\mathrm{Ca}_2$ both equal 1. The barrier heights between the wells are:

$$E_{\mathrm{out},A} = 2$$
$$E_{A,B} = 2$$
$$E_{B,C} = 6 - 1\,\mathrm{Ca}_1 - 1.5\,\mathrm{Ca}_2$$
$$E_{\mathrm{C,in}} = 6.5 - 1\,\mathrm{Ca}_1 - 2\,\mathrm{Ca}_2$$

The current is then:

$$I = -\eta\left(k_{21}\left(AXX + AXC + ABX\right) - k_{12}C_{\mathrm{out}}\left(XXX + XBX + XXC\right)\right) \tag{1}$$

where $\eta$ is the conversion from ions per microsecond to picoAmps (pA): 1 pA = 6.24159 ions/micro, so $eta = 1/6.24159$.

#### 2 A calcium binding model

We assume that the probability of identifying 1 or 2 $Ca^{2+}$ bound at each site is:

$$P_1 = \frac{[\mathrm{Ca}^{2+}]}{K_{d1} + [\mathrm{Ca}^{2+}] + [\mathrm{Ca}^{2+}]^2/K_{d2}} \tag{2}$$

$$P_2 = \frac{[\mathrm{Ca}^{2+}]^2}{K_{d1}K_{d2} + K_{d2}[\mathrm{Ca}^{2+}] + [\mathrm{Ca}^{2+}]^2}, \tag{3}$$

where $[Ca^{2+}]$ is the intracellular calcium concentration (in $\mu$M) and $K_{d1}$ and $K_{d2}$ are the binding constants for the first (upper) and second (lower) calcium ion binding to their respective locations. Here, $K_{d1} < K_{d2}$ indicating that the first binding event is stronger, and each has a voltage dependence of binding:

$$K_{d1} = \exp\bigl(E_1 - 4\varepsilon_1 V_r\bigr) \quad \text{and} \quad K_{d2} = \exp\bigl(E_2 - 4\varepsilon_2 V_r\bigr) \tag{4}$$

where $E_{1/2}$ are the voltage independent binding energy in units of $kT$ and $2\varepsilon_{1/2}$ are the fraction of the membrane electric field that each divalent cation moves through the field to reach the binding site, which is doubled since this is a divalent ion. Calculated from our electrostatics calculations (see Table S5). This model is the equilibrium solution to a sequential kinetic model of binding.

**Table S3. Membrane potential profile across Cluster 12 medoid structure.** Electrostatic potential at specific ion sites determined from Non-linear Poisson Boltzmann calculations using the parameters in Table 1, no protein charges, and a reduced membrane potential $V_r = -1$.

| | |
|---|---|
| -0.07 | $Cl^-$ site A |
| -0.14 | $Cl^-$ site B |
| -0.76 | $Cl^-$ site C |
| -0.84 | $Ca^{2+}$ upper site / first binder |
| -0.96 | $Ca^{2+}$ lower site / second binder |

**Table S4. Model parameters for the seven-state, three-site kinetic model.** The $\varepsilon$ values were determined from electrostatic calculations, while the rate constants were determined by scaling the overall conductance to match experiment.

| | |
|---|---|
| $k'_0 = 1{,}000$ $M^{-1}$ $\mu s^{-1}$ | $k_0 = 1{,}000$ $\mu s^{-1}$ |
| $\varepsilon_{12} = -0.035$ | $\varepsilon_{21} = 0.035$ |
| $\varepsilon_{23} = -0.035$ | $\varepsilon_{32} = 0.035$ |
| $\varepsilon_{34} = -0.315$ | $\varepsilon_{43} = 0.315$ |
| $\varepsilon_{41} = -0.115$ | $\varepsilon_{14} = 0.115$ |

**Table S5. Model parameters calcium binding.**

| | | |
|---|---|---|
| $E_1 = 5.5\ kT$ | $\varepsilon_1 = 0.0825$ | $Ca^{2+}$ upper site / first binder |
| $E_2 = -4\ kT$ | $\varepsilon_2 = 0.022$ | $Ca^{2+}$ lower site / second binder |

### Additional Supplemental Information

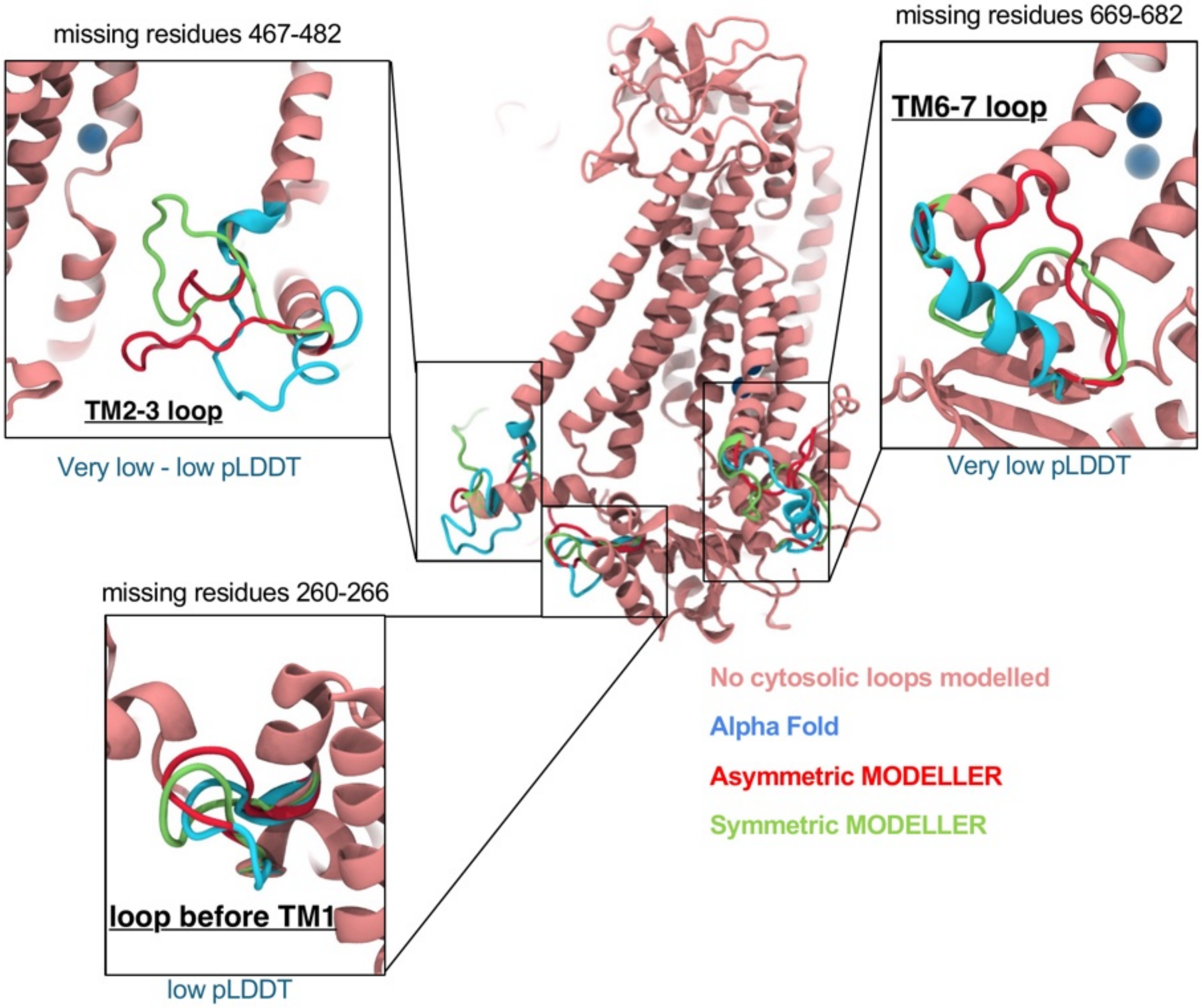


**Figure S1. Modeled missing loops in 1PBC/$Ca^{2+}$ -bound TMEM16A.** Computational models for three missing cytosolic loops (residues 260-266, 467-482, and 669-682) were predicted using MODELLER[1] (red and green) and AlphaFold[2] (blue). Qualitative descriptions of the AlphaFold per-residue confidence scores (pLDDT) are shown in the blue text. MODELLER-based loops in the symmetrized dimer are shown in red.

**Table S1. Summary of TMEM16A structures and pore characteristics in simulated and experimentally solved TMEM16A structures.** * Denotes conformations predicted by simulation. 5OYB* is a simulated structure from Jia and Chen[3]. All structures are of the 'ac' mTMEM16A variant except for 6BGJ and 6BGI which are of the 'a' variant. Distances and radii calculations ignore hydrogen atoms in simulated structures. Minimum pore radii only reported between site A and site C (see Fig. 2).

| structure/ conformation | Resolution (Å) | Minimum pore radius radius | L547-I641 Cα distance (Å) | # bound $Ca^{2+}$ | $PIP_2$ (Y/N) | inhibitor (Y/N) |
|---|---|---|---|---|---|---|
| 7ZK3* | N/A | 2.0 | 14.0 | 6 | N | N |
| 7ZK3* | N/A | 1.3 | 10.7 | 6 | N | N |
| 7ZK3* | N/A | 1.6 | 8.0 | 6 | N | N |
| 7ZK3* | N/A | 1.7 | 9.8 | 6 | N | N |
| 7ZK3* | N/A | 0.7 | 9.0 | 6 | N | N |
| 7ZK3 | 2.85 | 1.1 | 8.6 | 6 | N | Y |
| 5OYB* | N/A | 1.1 | 10.9 | 4 | Y | N |
| 5OYB | 3.75 | 1.2 | 8.2 | 4 | N | N |
| 5OYG | 4.06 | 0.7 | 7.3 | 0 | N | N |
| 8QZC | 3.29 | 1.2 | 6.4 | 4 | N | N |
| 5NL2 | 6.6 | 2.1 | 7.1 | 0 | N | N |
| 7B5E I551A | 4.1 | 1.0 | 8.1 | 4 | Y | N |
| 7B5D I551A | 3.3 | 0.8 | 7.8 | 0 | Y | N |
| 7B5C | 3.7 | 0.4 | 7.8 | 4 | N | N |
| 6BGJ (a) | 3.8 | 1.2 | 7.6 | 2 | N | N |
| 6BGI (a) | 3.8 | 1.5 | 7.8 | 4 | N | N |

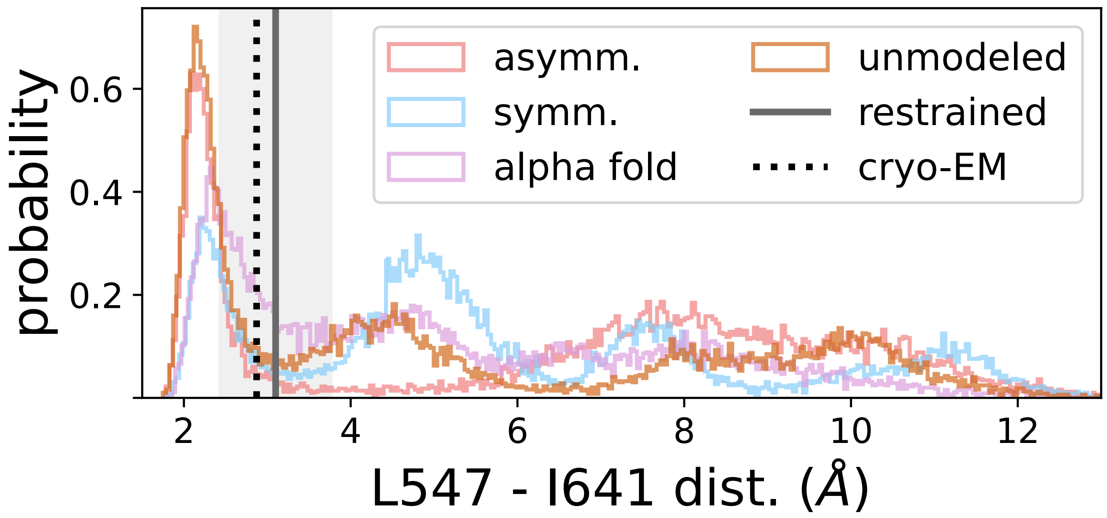


**Figure S2. Hydrophobic gate distances from 1PBC-removed simulations of each starting model.** Histograms of minimum heavy atom distances between L547 and I641 from aggregate simulation data (3 replicates, 1 μs long) for each TMEM16A loop model in Fig. S1: Asymmetric MODELLER (pink), symmetric MODELLER (blue), AlphaFold2 (purple), and no cytosolic loops (brown). The minimum distance in the cryo-EM structure is indicated by the black dotted line. Standard deviation and mean distances from a 1 kcal/mol·Å$^2$ backbone-restrained asymmetric model represented by the gray line and shaded region, respectively. Analysis excludes first 500 ns.

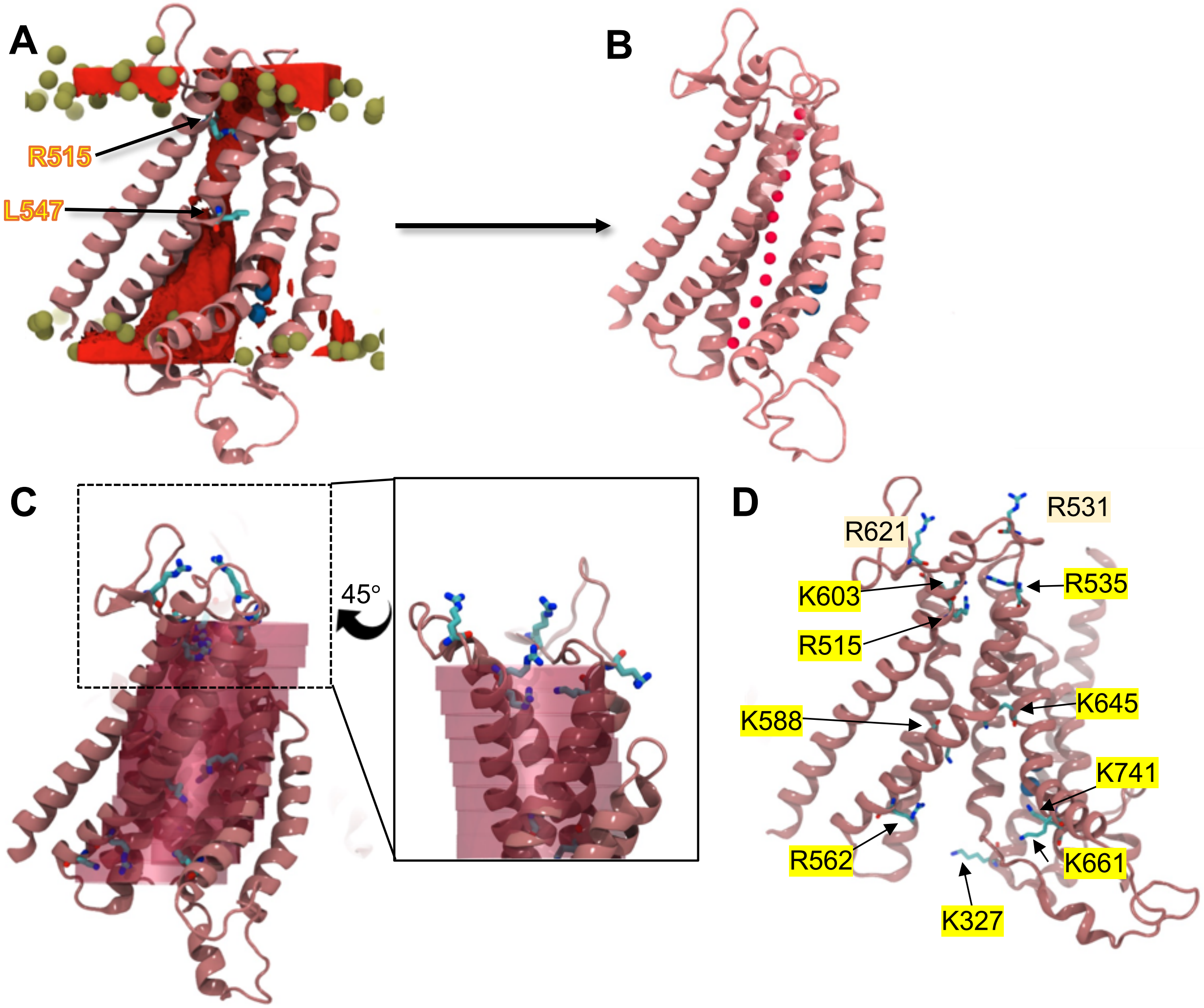


**Figure S3. Path coordinate for tracking ions and water in the TMEM16A pore**. **(A)** Snapshot from TMEM16A simulation with 1PBC removed showing the 3D water density (red). Only TM3-8 shown with lipid phosphate atoms (olive green). **(B)** Red beads mark the density-weighted mean of water coordinates in the pore every 2.5 Å in z (see Methods for details). **(C)** Cylinders used to calculate the water pathway in panel B. Each segment has a 12 Å diameter and 3.5 Å height. Each region was used as a search volume for identifying water and ions in the pore. **(D)** Basic residues in or near the pore.

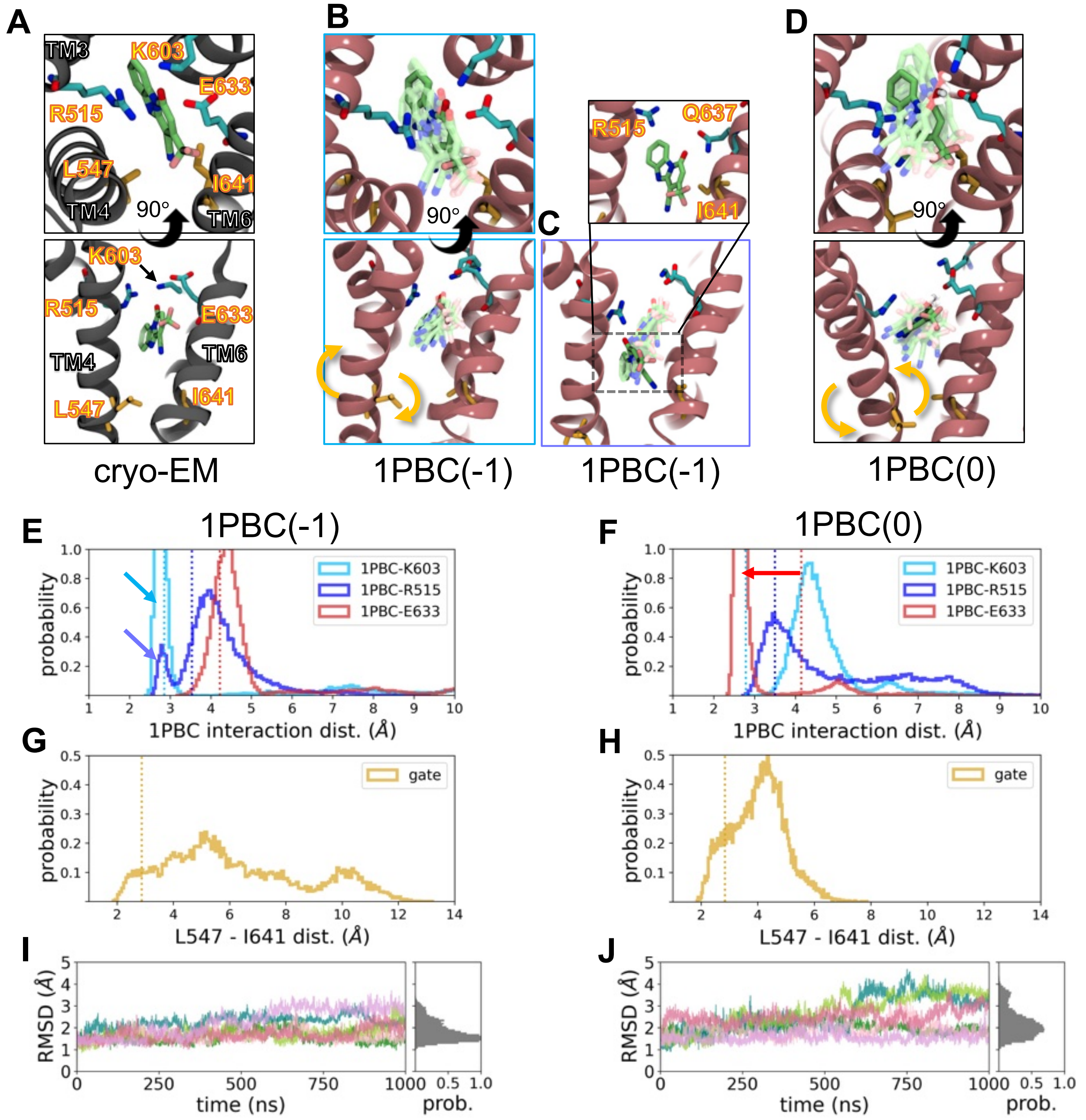


**Figure S4. TMEM16A pore dilates even when bound to inhibitor. (A)** Images of 1PBC/$Ca^{2+}$-bound TMEM16A (PDB ID 7ZK3) viewed from the extracellular space (top) and parallel to the plain of the membrane (bottom). **(B-D)** Snapshots from simulations of TMEM16A initiated from PDB ID 7ZK3 bound to anionic 1PBC (1PBC(-1)) (B, C) and protonated neutral 1PBC (1PBC(0)) (D). Snapshots taken from the same perspectives as in panel A. Positions of 1PBC from multiple evenly spaced frames are overlayed in the same image. Yellow arrows highlight the movement of TM4 from the starting cryo-EM position. **(E, F)** Histograms of distances between the 1PBC hydroxyl oxygen and nearby residues and **(G, H)** between L547 and I641 from aggregate simulation data for 1PBC(-1) (E, G) or 1PBC(0) (F, H). Blue and purple arrows highlight 1PBC(-1) distances sampled in snapshots in panel B outlined in the same color. **(I, J)** Cα RMSD of the extracellular-facing half of the ion pore defined by TM3-6 plotted over time and normalized probability of RMSD values for 1PBC(-1) (I) and 1PBC(0) (H). Analysis excludes first 500 ns. Simulations were initiated with symmetric MODELLER-based cytosolic loops.

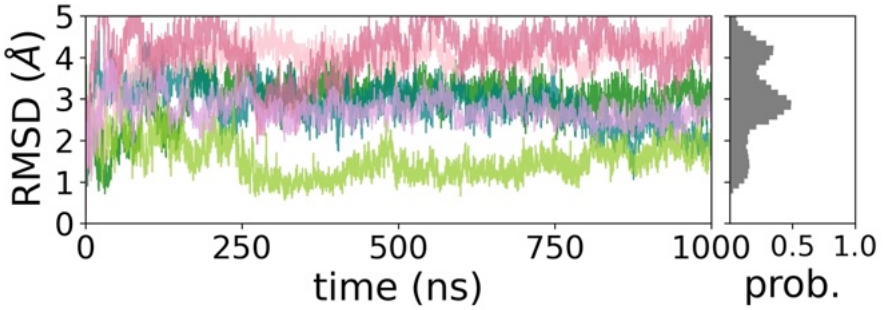


**Figure S5. TMEM16A pore dynamics after 1PBC removal.** C$\alpha$ RMSD of the extracellular-facing half of the pore in 3 replicates (one trace per subunit) of TMEM16A with 1PBC removed. Normalized probability density of RMSD values shown on the right in gray for aggregate simulation data. Simulations were initiated with symmetric MODELLER-based cytosolic loops.

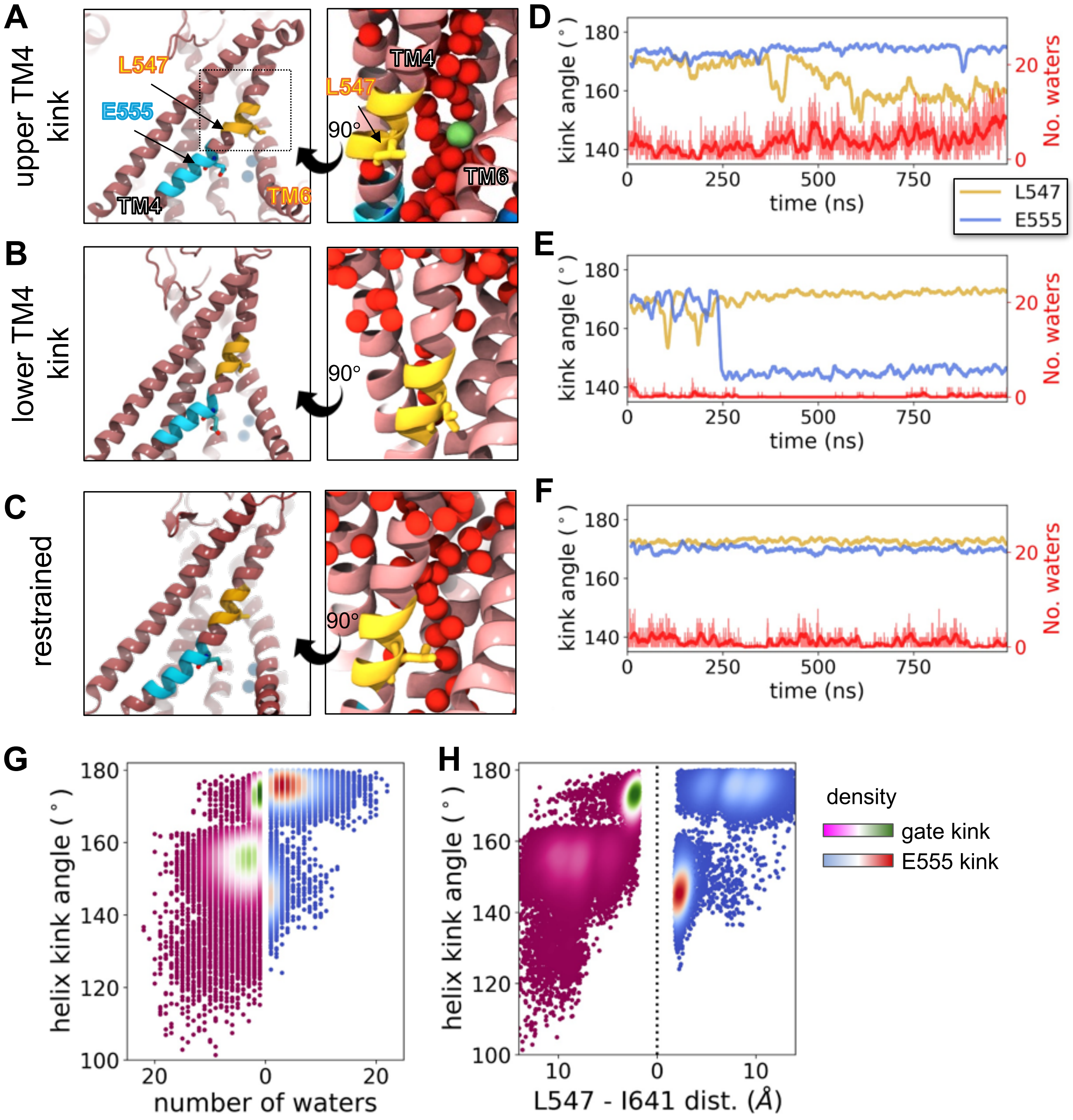


**Figure S6. Changes in TMEM16A pore hydration depend on kink location in TM4. (A-C)** Snapshots from unrestrained simulations with 1PBC removed in which TM4 kinks at the upper location L547 (A), lower location E555 (B), or remains straight in the restrained simulation (C). Enlarged view of each image on right show water oxygens (red spheres) and $Cl^-$ (green spheres). **(D-F)** Kink angle at L547 and E555 (left y-axis) and water flux (number of waters/0.25 ns passing L547, right y-axis) for the simulations in panels A-C, respectively. Kink angles plotted with a 2.5 ns rolling window average. **(G)** The total number of waters every 0.25 ns passing L547 plotted against the upper TM4 kink angle at L547 (pink/white/green values on left side of graph) or the lower angle at E555 (blue/white/red values on right side). **(H)** Minimum distance between hydrophobic gating residues L547 and I641 plotted against the upper kink angle at L547 (left values) or lower kink angle at E555 (right values). Points in panels G and H are colored by density: purple (least) to green (most) for the upper kink and blue (least) to red (most) lower kink.

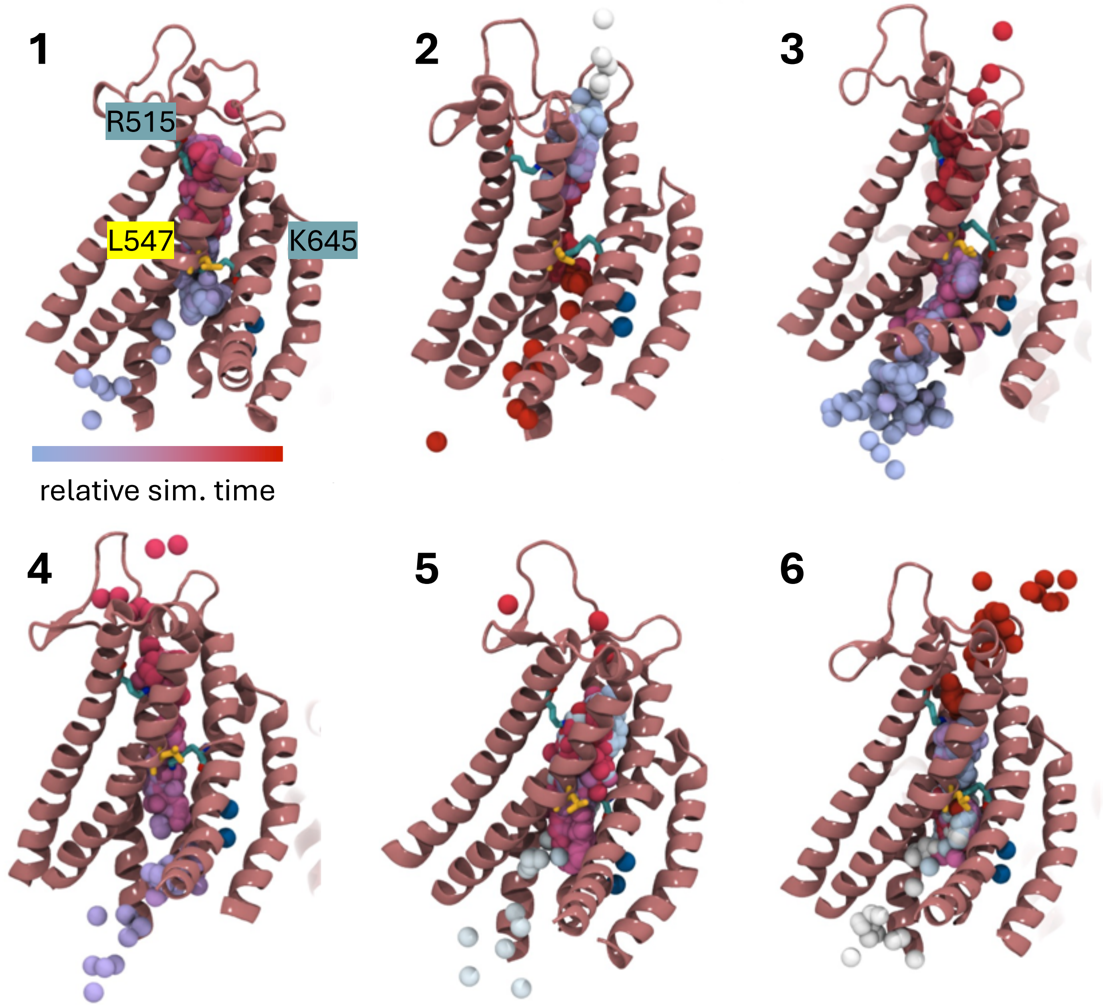


**Figure S7. Simulations of 1PBC-bound mTMEM16A with 1PBC removed samples six spontaneous full chloride permeation events.** Snapshots of permeating $Cl^-$ ions taken over the course of the event. Ion snapshots are colored by the relative simulation time for each individual event (blue - start, red - end). Protein coordinates correspond to the end of the permeation event. Permeation was observed in all four starting models: events 1 and 3 - asymmetric MODELLER loops; event 2 - asymmetric MODELLER loops; events 4 and 5 - unmodeled loops; and event 6 - AlphaFold2 loops.

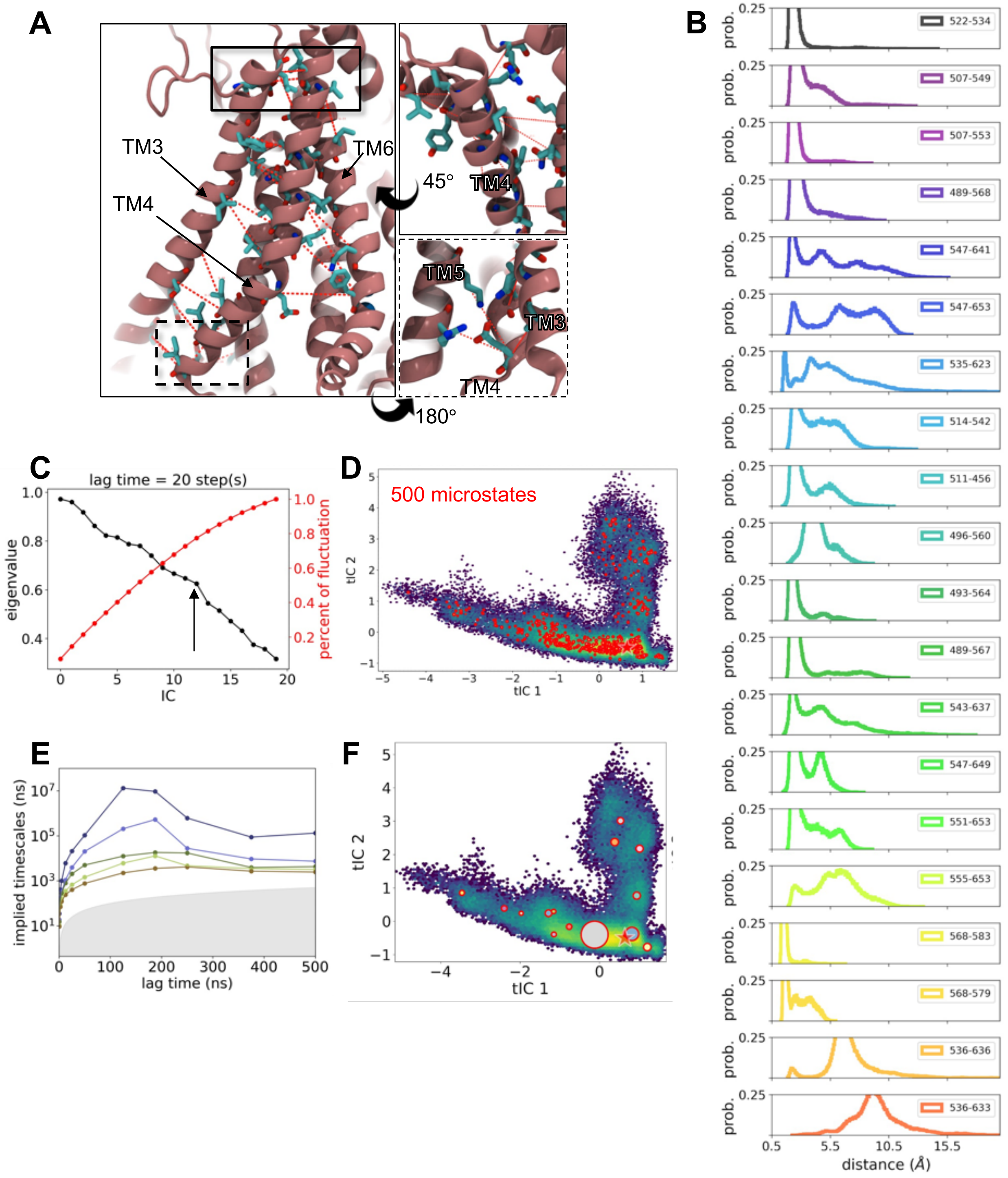


**Figure S8. Time Independent Component Analysis (tICA) of aggregate simulation data with 1PBC-removed. (A)** Twenty Cα-Cα distances values (red dotted lines) used as input for tICA. **(B)** Normalized probability distributions of all residue pair distances. **(C)** The eigenvalues (black, left axis) and cumulative percentage of the total fluctuation at each independent component (red, right axis). **(D)** Aggregate simulation data projected onto the first two tICs. Red dots represent cluster centers from MiniBatch K-means clustering on first 13 tICs. **(E)** Implied time scales (slowest 5) at different lag times from Markov state model (MSM) constructed from 500 microstates. tICA lag time (20 frames) and dimensions (13) that produced the highest MSM GMRQ score were used. The gray shaded region indicates times faster than the lag time. **(F)** Same data projection as in D with macrostate (cluster) centers calculated from MSM with a lag time of 18.75 ns (lowest MSM function value) using the PCC++ clustering method. The size of each macrostate center is proportional to the state population size. The red star indicates the starting cryo-EM position in the projected landscape. The projected simulation data is colored by log density (low density in purple to high density in yellow).

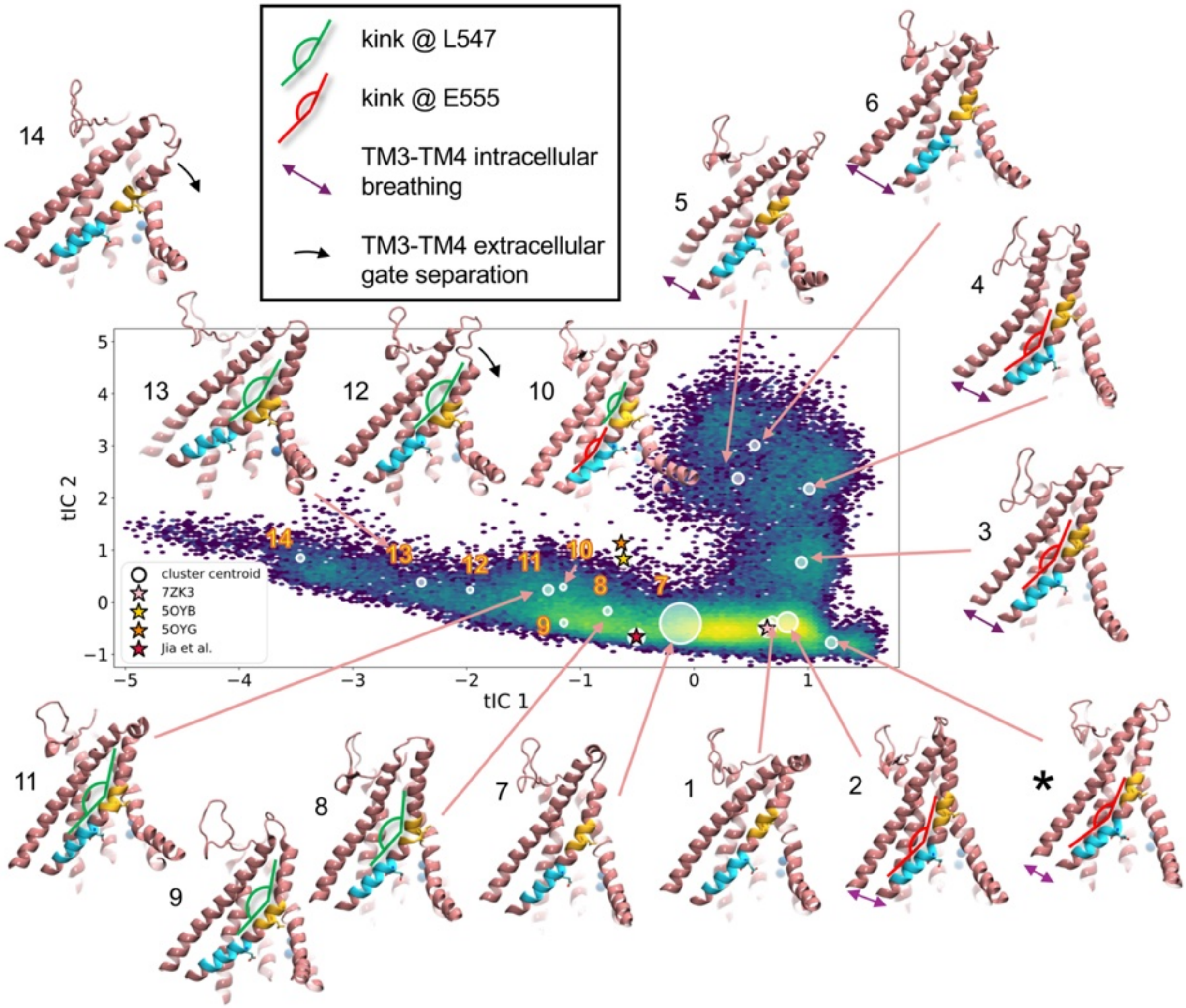


**Figure S9. Structural features of 15 clusters identified with tICA and MSM.** Aggregate simulation data projected onto the first two tICs with cluster centers and TMEM16A subunit structures representing the macrostate medoids. Only TM3-8 are shown. The cluster indicated by * was only sampled by simulations initiated with symmetric MODELLER-based cytosolic loops. Structural features of snapshots are highlighted on each structure: intracellular TM3-to-TM4 distance (purple double arrow), lower TM4 kink centered at E555 (red angle), upper TM4 kink centered at L547 (green angle), and TM4 tilt into the membrane (black curved arrow). The upper portion of TM4 is yellow and the lower portion is cyan.

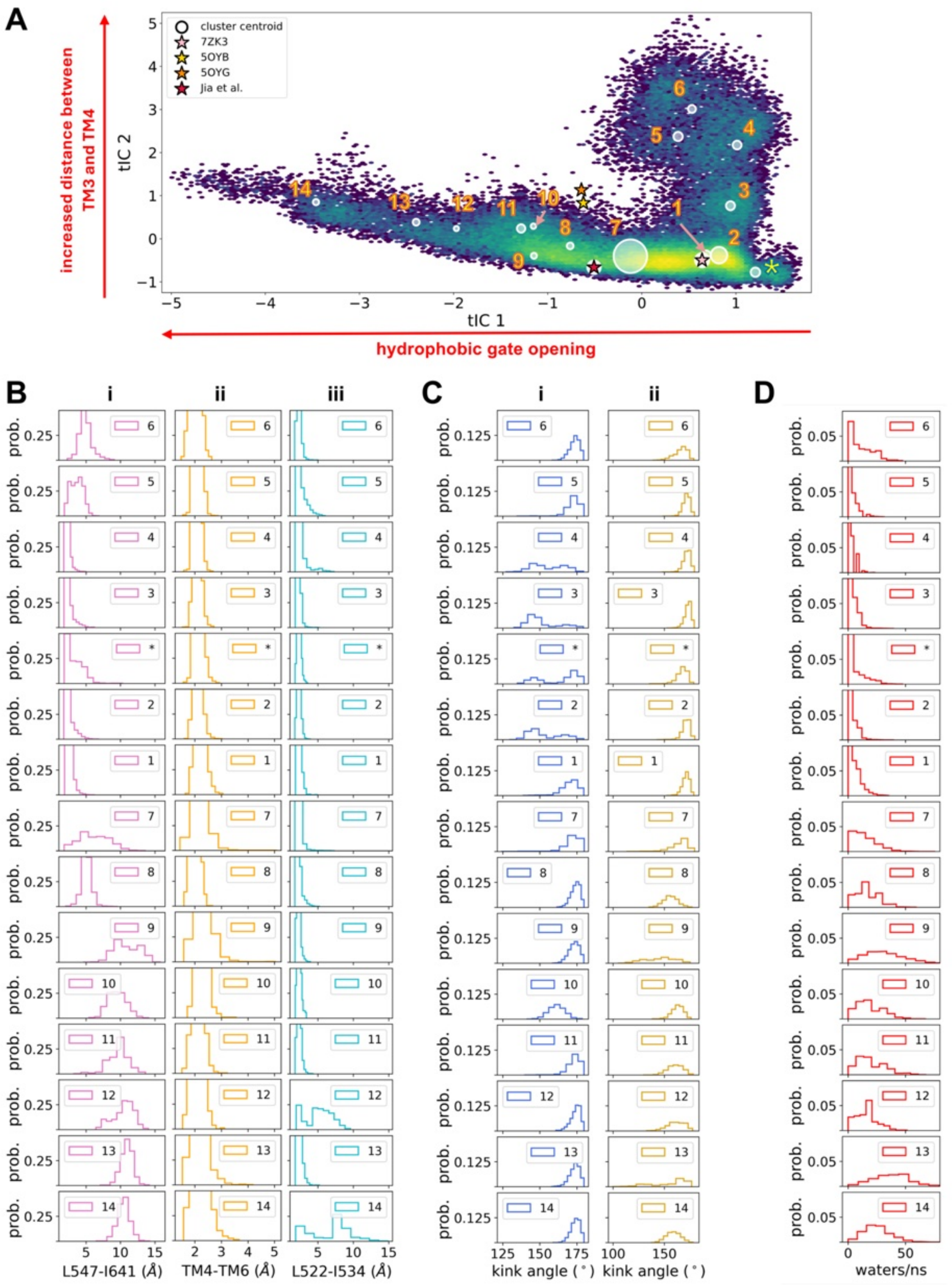


**Figure S10. Structural and functional properties of 15 clusters identified with tICA and MSM. (A)** Aggregate simulation data projected onto the first two tICs with cluster centroids calculated from the MSM (see Fig. S8-S9 for full description). Normalized probability distributions of data collected from simulation frames in each cluster: **(B)** minimum heavy atom sidechain distance between residues: (i) L547 and I641, (ii) 543-551 (TM4) and 641-653 (TM6), and (iii) L522 and I534, **(C)** TM4 kink angles centered on: (i) E555 and (ii) L547, and **(D)** number of waters per ns passing L547 from the intracellular bath.

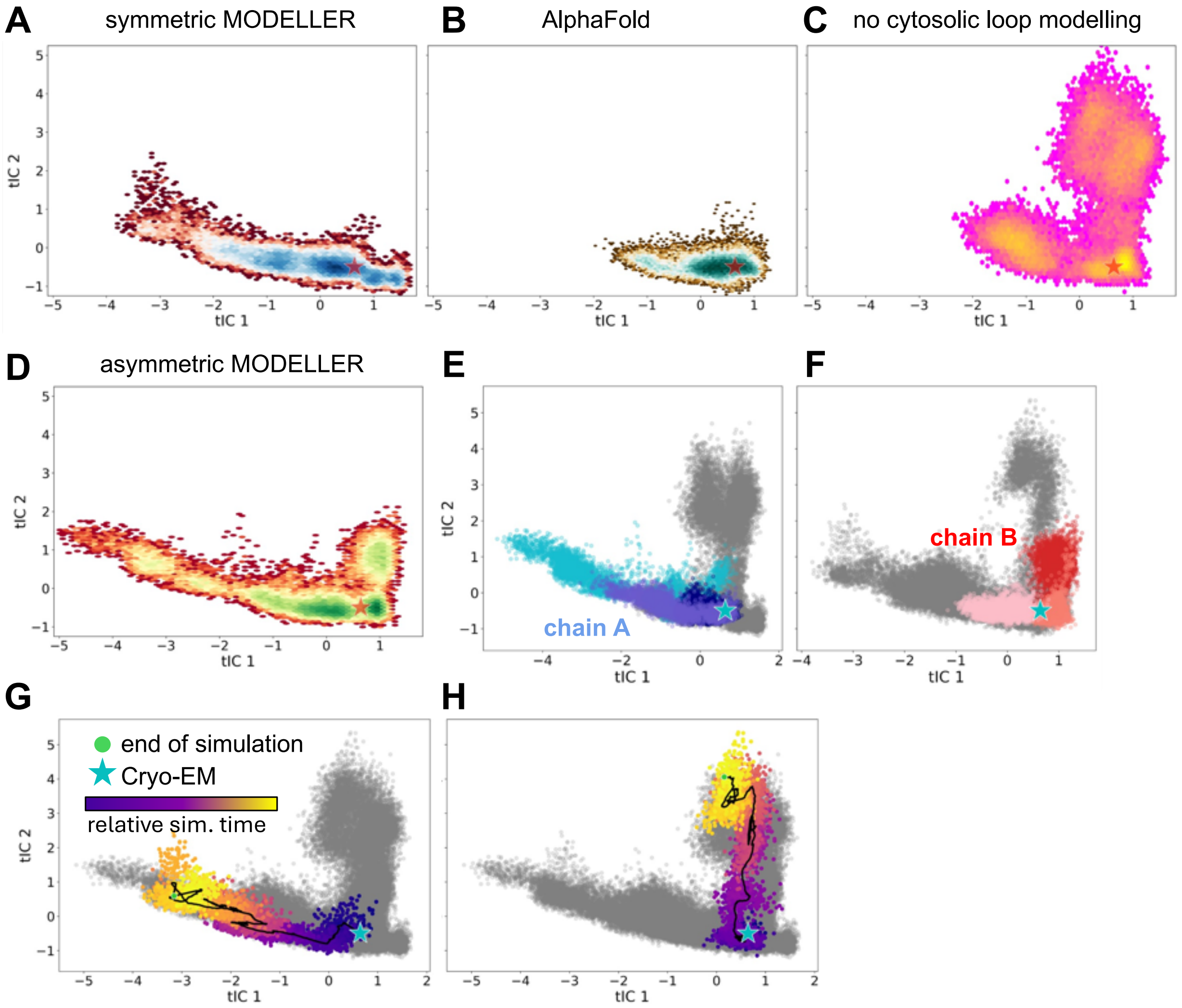


**Figure S11. Simulations of TMEM16A with different cytosolic loops models sample common and diverging regions of tIC space. (A-D)** Aggregate simulation data from each class of models projected onto tIC 1 and 2: (A) symmetric MODLLER, (B) AlphaFold2, (C) no loops, and (D) asymmetric MODELLER. Data colored by log of density with the high density dark blue, teal, yellow, and green for data in A-D, respectively. The initial cryo-EM structure is the star in all panels. **(E, F)** All simulation data (gray) shown separately for chain A (E) and chain B (F). As an example, the colored data in both panels is from simulations initiated with asymmetric MODELLER-based loops. Chain A of the asymmetric model was used to build the symmetric model (panel A), which may explain similarities in panels A and D, but stochastic differences cannot be ruled out. **(G, H)** Simulation data from all models projected on the first two tICs (gray) overlayed with a single time series trajectory of a model built with symmetric loops in MODELLER (G) and one built without loops (H). These trajectories progress from dark blue, at the cryo-EM structure (sky blue star), to yellow ending at the green dot.

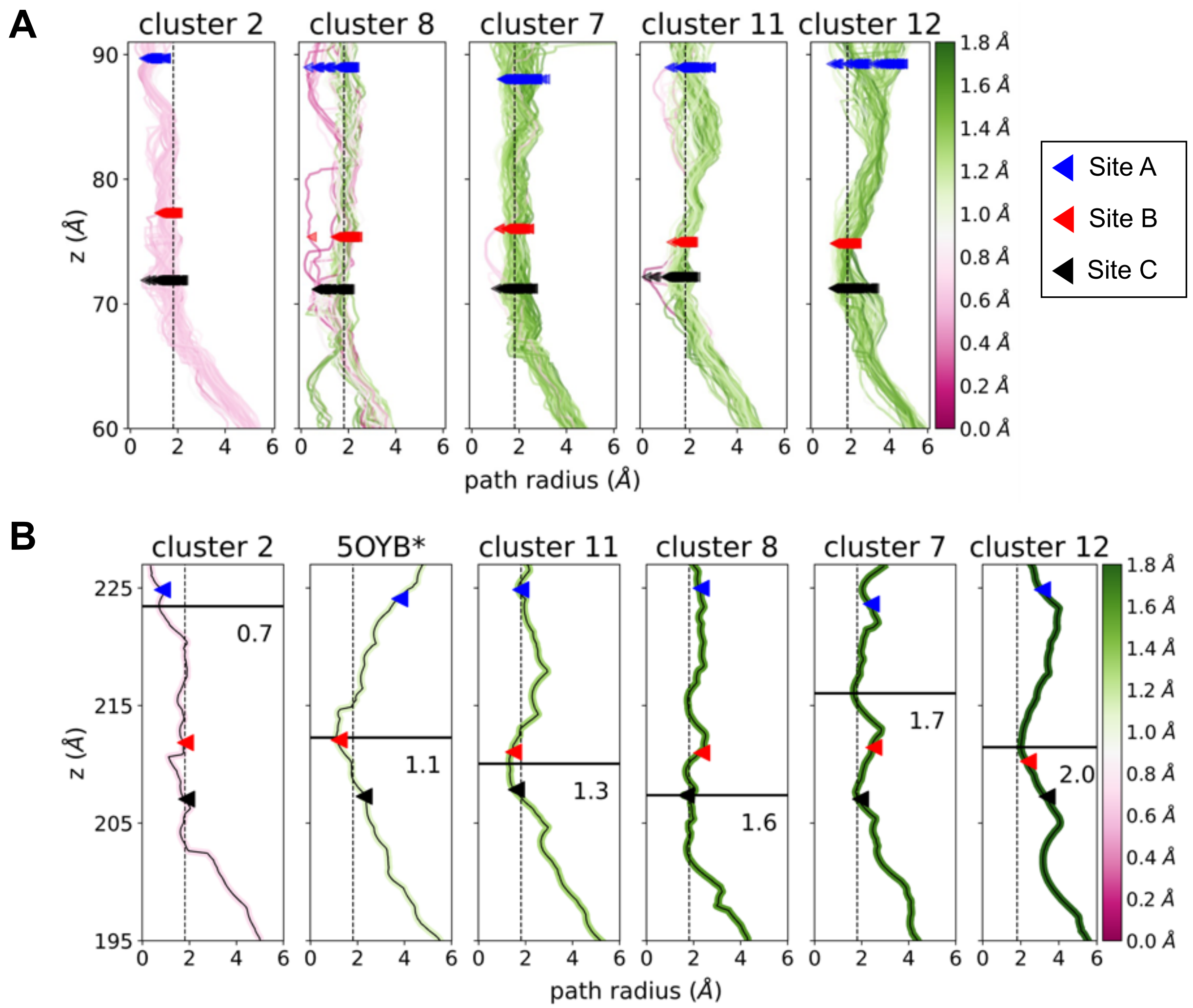


**Figure S12. Pore radii calculated for select clusters. (A)** Restrained simulations were initiated from the medoid of specific clusters (1 kcal/mol·$Å^2$ backbone-restraints on TM3-8), and the pore radii for each snapshot were calculated with HOLE2[4]. Conformations were taken every 12.5 ns, and the color of each curve corresponds to the minimum value according the heat map on the right. Triangles mark the centroid z-positions of the residues forming the $Cl^-$ interaction sites: R515/K603 (site A), R515/K645 (site B), and K645/K555 (site C). **(B)** Average pore radii calculated from the data in panel A. The second panel is the profile for the predicted open state from Jia and Chen (5OYB*)[3]. Black horizonal bars mark the location of the minimum pore radius with the value given in Ångstroms. All other features are as in panel A.

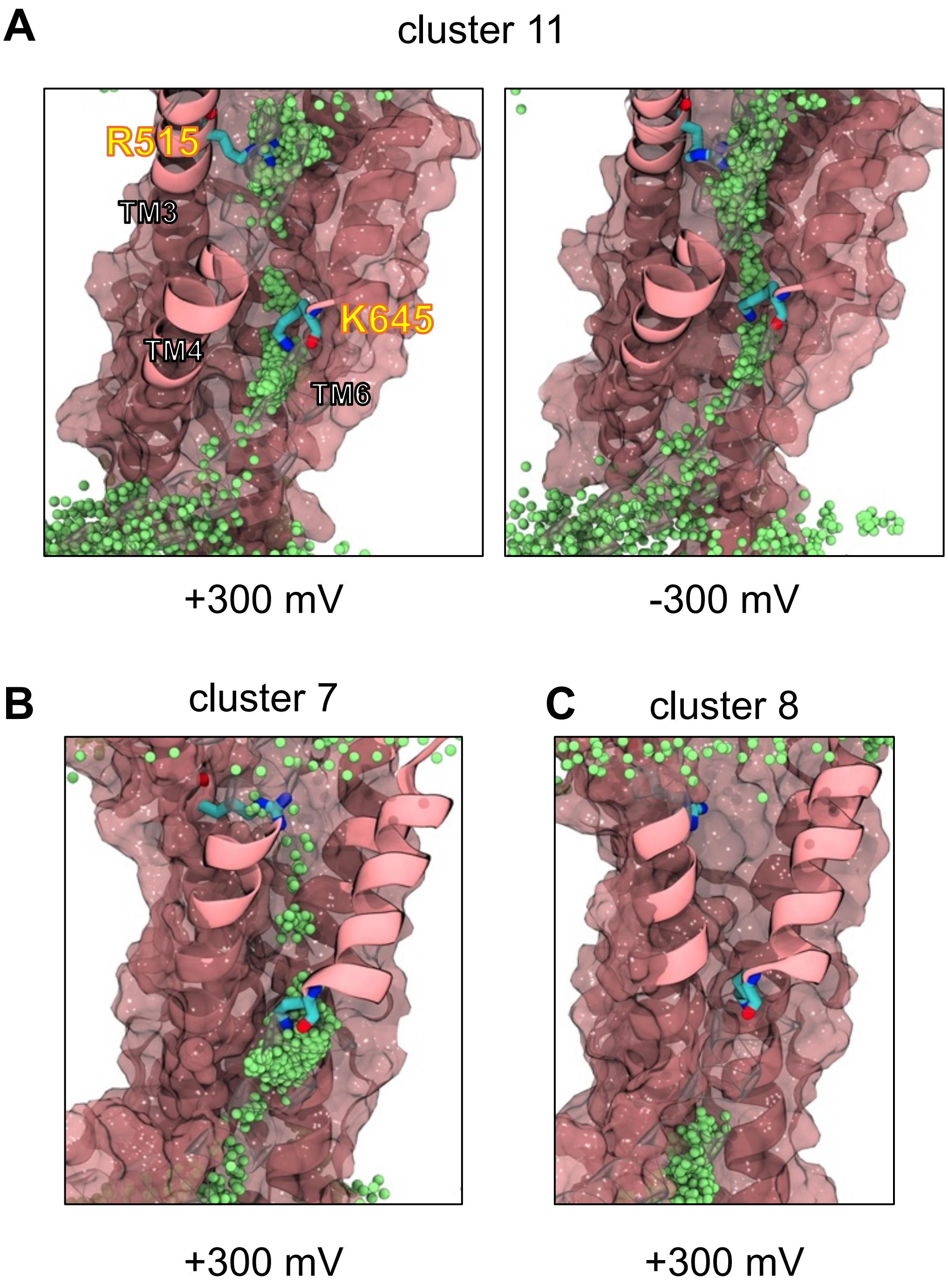


**Figure S13. Chloride positions in TMEM16A pore. (A)** Representative snapshots taken from TMEM16A simulation initiated from cluster 11 medoid with -300 mV (left) or 300 mV (right) applied voltage. Only TM3-6 are shown (pink) with overlays of $Cl^-$ every 0.5 ns (green spheres). **(B)** Snapshot from simulation initiated from cluster 7 with 300 mV applied potential. **(C)** Snapshot from simulation initiated from cluster 8 with 300 mV applied potential. See Fig. S9-S10 for full cluster descriptions.

**Table S2. Number of chloride permeations events and maximum conductance in applied voltage simulations.** Clusters 2 and 8 revealed no permeation events across the range of voltages.

| Cluster 7 | | Cluster 11 | | Cluster 12 | | |
|---|---|---|---|---|---|---|
| **Voltage (mV)** | **Number of Events** | **Voltage (mV)** | **Number of Events** | **Voltage (mV)** | **Number of Events** | **I (pA) with 95% CI range** |
| -300 | 0 | -300 | 2 | -350 | 8 | -1.3 (-2.3, -0.8) |
| -250 | 0 | -250 | 1 | -285 | 6 | 1.0 (-1.9, -0.5) |
| -200 | 0 | -200 | 0 | -230 | 6 | 1.0 (-1.9, -0.5) |
| -150 | 0 | -150 | 2 | -170 | 2 | -0.3 (-1.0, -0.1) |
| 0 | 0 | 0 | 0 | 0 | 1 | 0.2 (0.1, 0.7) |
| +150 | 0 | +150 | 0 | +170 | 3 | 0.5 (0.2, 1.2) |
| +200 | 1 | +200 | 0 | +230 | 7 | 1.1 (0.6, 2.1) |
| +250 | 0 | +250 | 0 | +285 | 2 | 0.3 (0.1, 1.0) |
| +300 | 1 | +300 | 3 | +350 | 20 | 3.2 (2.3, 4.7) |
| Total events: | 2 | Total events: | 8 | Total events: | 55 | -- |

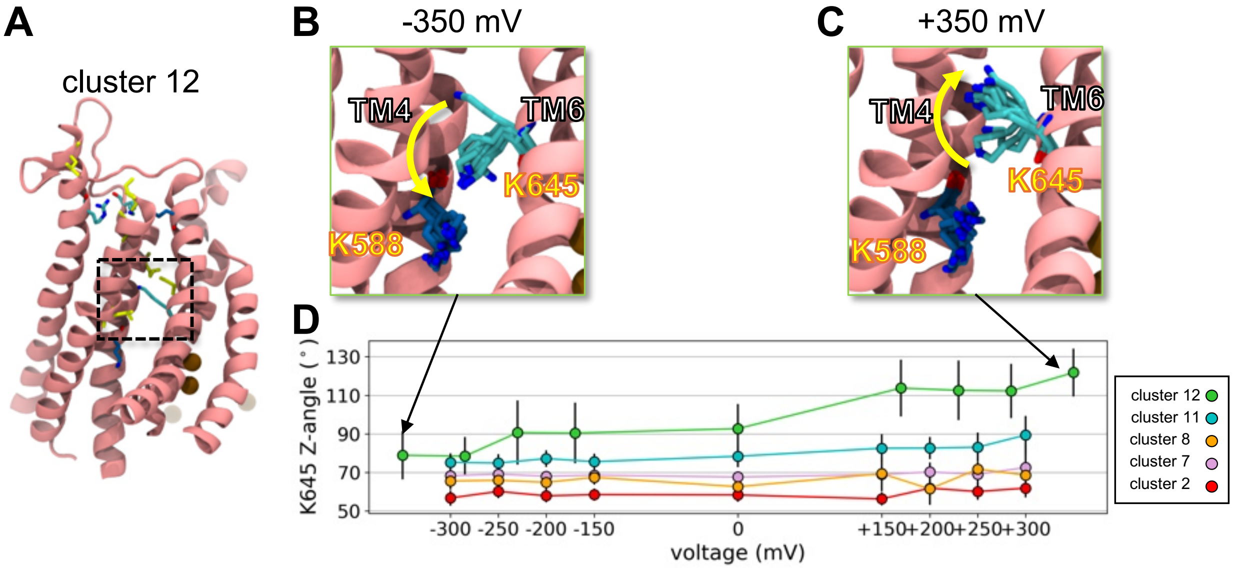


**Figure S14. K645 conformation is voltage dependent. (A)** Snapshot from simulations initiated from cluster 12 medoid under 350 mV applied voltage. **(B, C)** Cartoon representation of the inner vestibule $Cl^-$ interaction site (site C in Fig. 2B) with overlay of the K645 and K588 sidechains every 50 ns under -350 mV (panel B) and 350 mV voltage (panel C). **(D)** The average angle between the K645 sidechain principal axis and the z unit vector for simulations initiated from medoids of 5 different clusters. Black bars indicate the standard deviation of each angle.

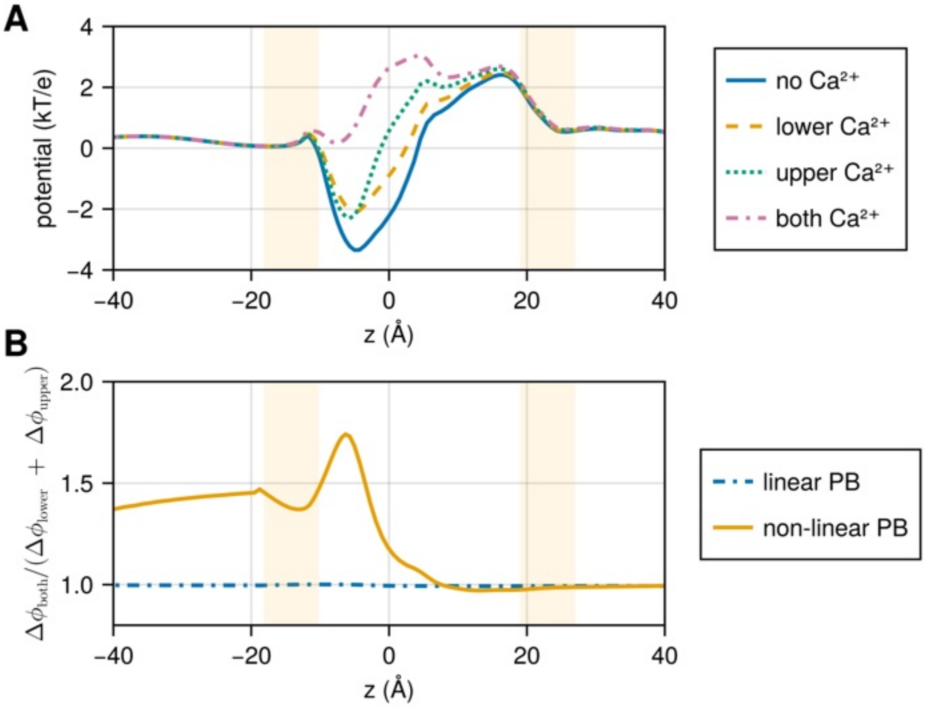


**Figure S15. Influence of $Ca^{2+}$ on the electrostatic profile computed with continuum electrostatics using the cluster 12 medoid. (A)** Potential along the $Cl^-$ permeation pathway, from the cytoplasmic side on the left to the extracellular side on the right, due to TMEM16A atomic partial charges and $Ca^{2+}$ ions in sites near the pathway. Solid blue: no $Ca^{2+}$ ions present. Dotted green: a $Ca^{2+}$ in the upper/1$^{st}$ binding site only. Dashed yellow: a $Ca^{2+}$ in the lower/2$^{nd}$ binding site only. Dash-dotted purple: $Ca^{2+}$ in both sites. The vertical yellow bands mark the z positions of the hydrophilic headgroups of a nominally undistorted lipid bilayer. **(B)** The two $Ca^{2+}$ ions act cooperatively to reduce the barrier for $Cl^-$ entry from the cytoplasm as captured by non-linear Poisson-Boltzmann calculations. The ratio plotted on the y-axis compares the change in potential caused by adding two $Ca^{2+}$ ions simultaneously ($\Delta\phi_{both}$) to the sum of the potential changes resulting from adding a single $Ca^{2+}$ ion to site one and to site two individually ($\Delta\phi_{lower} + \Delta\phi_{upper}$). The linear calculation (blue dash-dotted line) is 1.0 across the entire permeation pathway since the fields add linearly at this level of theory, but the non-linear calculation shows significant deviation from linear behavior in the lower vestibule and permeation pathway (yellow curve) resulting in enhanced $Cl^-$ stabilization at site C due to the addition of the second $Ca^{2+}$.

**Video S1. Simulation of simulated open TMEM16A state (cluster 12) with 230 mV applied membrane potential.** Video includes the first 500 ns of the full 1 μs simulation. Only chloride ions (colored spheres) within 10 Å of the protein are shown as well as bound $Ca^{2+}$ ions (blue spheres). Basic pore lining residues K515, R353, K602, K645, and K555 are shown as cyan sticks. Only TM3-8 protein backbone shown. Simulations were run with 1 kcal/mol·$Å^2$ backbone-restraints on TM3-8.

**Video S2. Simulation of simulated open TMEM16A state (cluster 12) with -230 mV applied membrane potential.** Video includes the first 500 ns of the full 1 μs simulation. Only chloride ions (colored spheres) within 10 Å of the protein are shown as well as bound $Ca^{2+}$ ions (blue spheres). Basic pore lining residues K515, R353, K602, K645, and K555 are shown as cyan sticks. Only TM3-8 protein backbone shown. Simulations were run with 1 kcal/mol·$Å^2$ backbone-restraints on TM3-8.

**Kinetic Model S1.** *Stephens_TMEM16A_currents.mmd.* Mathematical model used to calculate currents in Figure 6. This model can be run by downloading the Berkeley Madonna software from:

`https://berkeley-madonna.myshopify.com/pages/download`

and running in the free demo mode.

#### SUPPLEMENTAL REFERENCES


1. Fiser, A., & Sali, A. (2003). ModLoop: Automated modeling of loops in protein structures. *Bioinformatics*. 19, 2500–2501.
2. Jumper, J., Evans, R., Pritzel, A., Green, T., Figurnov, M., Ronneberger, O., Tunyasuvunakool, K., Bates, R., Žídek, A., Potapenko, A., Bridgland, A., Meyer, C., Kohl, S.A.A., Ballard, A.J., Cowie, A., Romera-Paredes, B., Nikolov, S., Jain, R., Adler, J., … Hassabis, D. (2021). Highly accurate protein structure prediction with AlphaFold. *Nature*. 596, 583–589.
3. Jia, Z., & Chen, J. (2021). Specific PIP2 binding promotes calcium activation of TMEM16A chloride channels. *Communications Biology*. 4, 90.
4. Smart, O.S., Neduvelil, J.G., Wang, X., Wallace, B.A., & Sansom, M.S.P. (1996). HOLE: A program for the analysis of the pore dimensions of ion channel structural models. *Journal of Molecular Graphics*. 14, 354–360.
